# A genetically-encoded fluorescent sensor enables rapid and specific detection of dopamine in flies, fish, and mice

**DOI:** 10.1101/332528

**Authors:** Fangmiao Sun, Jianzhi Zeng, Miao Jing, Jingheng Zhou, Jiesi Feng, Scott F. Owen, Yichen Luo, Funing Li, Takashi Yamaguchi, Zihao Yong, Yijing Gao, Wanling Peng, Lizhao Wang, Siyu Zhang, Jiulin Du, Dayu Lin, Min Xu, Anatol C. Kreitzer, Guohong Cui, Yulong Li

**Affiliations:** State Key Laboratory of Membrane Biology, Peking University School of Life Sciences, Beijing 100871, China; PKU-IDG/McGovern Institute for Brain Research, Beijing 100871, China; Peking-Tsinghua Center for Life Sciences, Beijing 100871, China; Laboratory of Integrative Neuroscience, National Institute on Alcohol Abuse and Alcoholism, National Institutes of Health, Rockville, MD, USA; Gladstone Institutes, San Francisco, CA 94158, USA; Institute of Neuroscience, State Key Laboratory of Neuroscience, CAS Center for Excellence in Brain Science and Intelligence Technology, Shanghai Institutes for Biological Sciences, Chinese Academy of Sciences, Shanghai, China; Neuroscience Institute, New York University School of Medicine, New York, NY, USA; College of Biological Sciences, China Agricultural University, Beijing 100193, China; Chinese Academy of Sciences Center for Excellence in Brain Science and Intelligence Technology, Chinese Academy of Sciences, Shanghai 200031, China; Institute of Neuroscience, Chinese Academy of Sciences, Shanghai 200031, China; Shanghai Jiao Tong University School of Medicine, Shanghai 200025, China; University of Chinese Academy of Sciences, Beijing, China; Department of Psychiatry, New York University School of Medicine, New York, NY, USA; Center for Neural Science, New York University, New York, NY, USA; Gladstone Institutes, San Francisco, CA, USA; Department of Neurology, UCSF, San Francisco, CA, USA; Kavli Institute for Fundamental Neuroscience, UCSF, San Francisco, CA, USA; UCSF Weill Institute for Neurosciences, UCSF, San Francisco, CA, USA; Department of Physiology, UCSF, San Francisco, CA 94158, USA

## Abstract

Dopamine (DA) is a central monoamine neurotransmitter involved in many physiological and pathological processes. A longstanding yet largely unmet goal is to measure DA changes reliably and specifically with high spatiotemporal precision, particularly in animals executing complex behaviors. Here we report the development of novel genetically-encoded GPCR-Activation-Based-DA (GRAB_DA_) sensors that enable these measurements. In response to extracellular DA rises, GRAB_DA_ sensors exhibit large fluorescence increases (ΔF/F_0_∼90%) with sub-second kinetics, nanomolar to sub-micromolar affinities, and excellent molecular specificity. Importantly, GRABDA sensors can resolve a single-electrical-stimulus evoked DA release in mouse brain slices, and detect endogenous DA release in the intact brains of flies, fish, and mice. In freely-behaving mice, GRABDA sensors readily report optogenetically-elicited nigrostriatal DA release and depict dynamic mesoaccumbens DA changes during Pavlovian conditioning or during sexual behaviors. Thus, GRAB_DA_ sensors enable spatiotemporal precise measurements of DA dynamics in a variety of model organisms while exhibiting complex behaviors.

## Introduction

Dopamine is a crucial monoamine neurotransmitter in both vertebrates and invertebrates. In the vertebrate central nervous system, DA regulates a wide range of complex processes, including reward signaling (Schultz, 2016; Wise, 2004), reinforcement learning (Holroyd and Coles, 2002), attention (Nieoullon, 2002), motor control (Graybiel et al., 1994), arousal (Kume et al., 2005; Wisor et al., 2001), and stress (Abercrombie et al., 1989). In the human brain, impaired DA transmission is associated with neuropsychiatric disorders and neurodegenerative diseases, including attention deficit hyperactivity disorder (Cook Jr et al., 1995), schizophrenia (Howes and Kapur, 2009) and Parkinson’s disease (Lotharius and Brundin, 2002). Moreover, psychostimulants, such as cocaine and amphetamine, exert their addictive effects by targeting components in the DA signaling pathway and by altering extracellular DA levels (Di Chiara and Imperato, 1988; Giros et al., 1996; Hernandez and Hoebel, 1988; Ritz et al., 1987).

Despite many important roles that DA plays in both physiological and pathological processes, precise measurements of the spatial and temporal patterns of DA release during complex behaviors are lacking. This gap in understanding due in large part to the limitation associated with existing methods for the real-time detection of endogenous DA release in the intact brain. Historically, intracerebral microdialysis has served as the gold standard for quantitative measurements of extracellular DA concentration. However, the relative slow sampling rate afforded by microdialysis is not well suited to detect dynamic changes in DA levels during complex and rapidly evolving behaviors, such as those that characterize social affiliation (Tidey and Miczek, 1996), aggression (van Erp and Miczek, 2000), and mating (Pfaus et al., 1990a, b). Fast-scan cyclic voltammetry (FSCV) is a temperamental electrochemical method that can measure changes in extracellular DA concentrations with 10-ms temporal resolution (Robinson et al., 2008; Rodeberg et al., 2017). However, because FSCV requires oxidization of DA molecules, it is difficult to distinguish DA from other structurally similar transmitters, such as norepinephrine (NE) (Robinson et al., 2003). Moreover, both microdialysis and FSCV require implantation of a relatively large probe or electrode (approximately 70-300 μm in diameter) into a specific brain region, which precludes the ability to obtain spatially precise measurements of endogenous DA release (Jaquins-Gerstl and Michael, 2015).

In lieu of direct measurements of extracellular DA, indirect methods, for example, measuring the activity of dopaminergic neurons (DANs) or the activation of DA receptor downstream targets, have also been used to approximate the dynamics of DA release. Genetic expression of a presynaptic tethered Ca^2+^ indicator in DANs has been used to indicate potential compartmentalized DA signals in the *Drosophila* olfactory pathway (Cohn et al., 2015). However, given the highly complex regulation of presynaptic Ca^2+^, the nonlinear relationship between intracellular Ca^2+^ and transmitter release, as well as the active uptake of extracellular DA by transporters, it is difficult to quantitatively translate Ca^2+^ signals precisely into extracellular DA levels. Cell-based DA reporters, such as CNiFERs (Muller et al., 2014), use transplated HEK293 cells constitutively expressing DA receptors together with an intracellular Ca^2+^ indicator to couple extracellular DA signals with the fluorescence increase. However, This approach requires cell transplantation, which may limit the broad usage of this method. Moreover, CNiFERs are not spatially sensitive to synaptically released DA and thus may be limited to measurements of volumetric transmission of DA. Finally, the TANGO assay and their next-generation versions (Barnea et al., 2008; Inagaki et al., 2012; Kim et al., 2017; Lee et al., 2017) have been used to measure endogenous DA release by coupling the β-arrestin signaling pathway to reporter gene expression. Although this approach enables the cell-type specific expression of the DA reporter and is suitable for *in vivo* measurements, the long signal amplification time (on the order of hours) required for this assay precludes the ability to monitor rapid, physiologically relevant dynamics of DA signaling in real time.

Here, we report the development of novel, genetically-encoded sensors that enable direct, rapid, sensitive, and cell-type specific detection of extracellular DA. These sensors, which we call GRAB_DA_ sensors, were engineered by coupling a conformationally sensitive circular-permutated EGFP (cpEGFP) to a selected human DA receptor. Through iterative engineering and optimization, we yielded two GRAB_DA_ sensors: GRAB_DA1m_, with medium DA affinity (EC_50_ ∼ 130 nM); and GRAB_DA1h_, with high DA affinity (EC_50_ ∼ 10 nM). We show that these two newly developed GRAB_DA_ sensors enable real-time detection of endogenous DA in acute brain slices of mice and in the intact brains of versatile animal models including flies, fish, and mice.

## Results

### Development and initial characterization of GRAB_DA_ sensors

To develop a genetically encoded sensor for DA, we sought to engineer naturally evolved DA receptors to couple a conformationally sensitive cpEGFP. We hypothesized that upon DA binding, the conformational changes in the receptor could alter the arrangement of the associated cpEGFP, resulting in a DA-dependent change in fluorescence. Indeed, a similar strategy was recently applied in creating the genetically encoded acetylcholine sensor GACh (Miao Jing et al., 2018), suggesting that this strategy could potentially be expanded to generate DA sensors.

We used a three-step approach to engineer and optimize GRAB_DA_ sensors (Fig. 1A-C): First, as the third intracellular loop (ICL3) of the G protein-coupled receptor (GPCR) links transmembrane V and VI that are thought to undergo large conformational changes upon ligand binding (Kruse et al., 2013), a cpEGFP was first inserted into the ICL3 of all five subtypes of human dopamine receptors (DRs); we subsequently focused on the D_2_R-cpEGFP chimera due to its superior membrane trafficking and relatively high affinity for DA (Beaulieu and Gainetdinov, 2011; Missale et al., 1998) (Fig. S1A,B). In the second step, the position of the cpEGFP insertion within the ICL3, and individual amino-acids in the linker region between cpEGFP and D_2_R were systematically screened (Fig. 1A,B). Finally, mutations at residues critical for receptors affinity for DA (Sung et al., 2016) were further introduced to expand the sensor’s response range (Fig. 1C). After screening a total of more than 432 variants, we chose two, GRAB_DA1m_ and GRAB_DA1h_, both of which have a ∼90% maximal ΔF/F_0_ response to the application of saturating levels of DA (Fig. 1B,E), but differ by an order of magnitude with respect to affinity for DA (130 nM for GRAB_DA1m_ and 10 nM for GRAB_DA1h_) (Fig. 1C). We also generated a corresponding mutant variant of each sensor (GRAB_DA1m-mut_ and GRAB_DA1h-mut_) by introducing C118A and S193N double mutations in the receptor’s putative DA-binding pocket in order to abolish the DA binding (Chien et al., 2010b; Wang et al., 2018) (Fig. S1D). These so-called “dead” mutants exhibited no DA-induced change in fluorescence compared with GRAB_DA1m_ and GRAB_DA1h_ (Fig. 1D).

**Figure 1:**
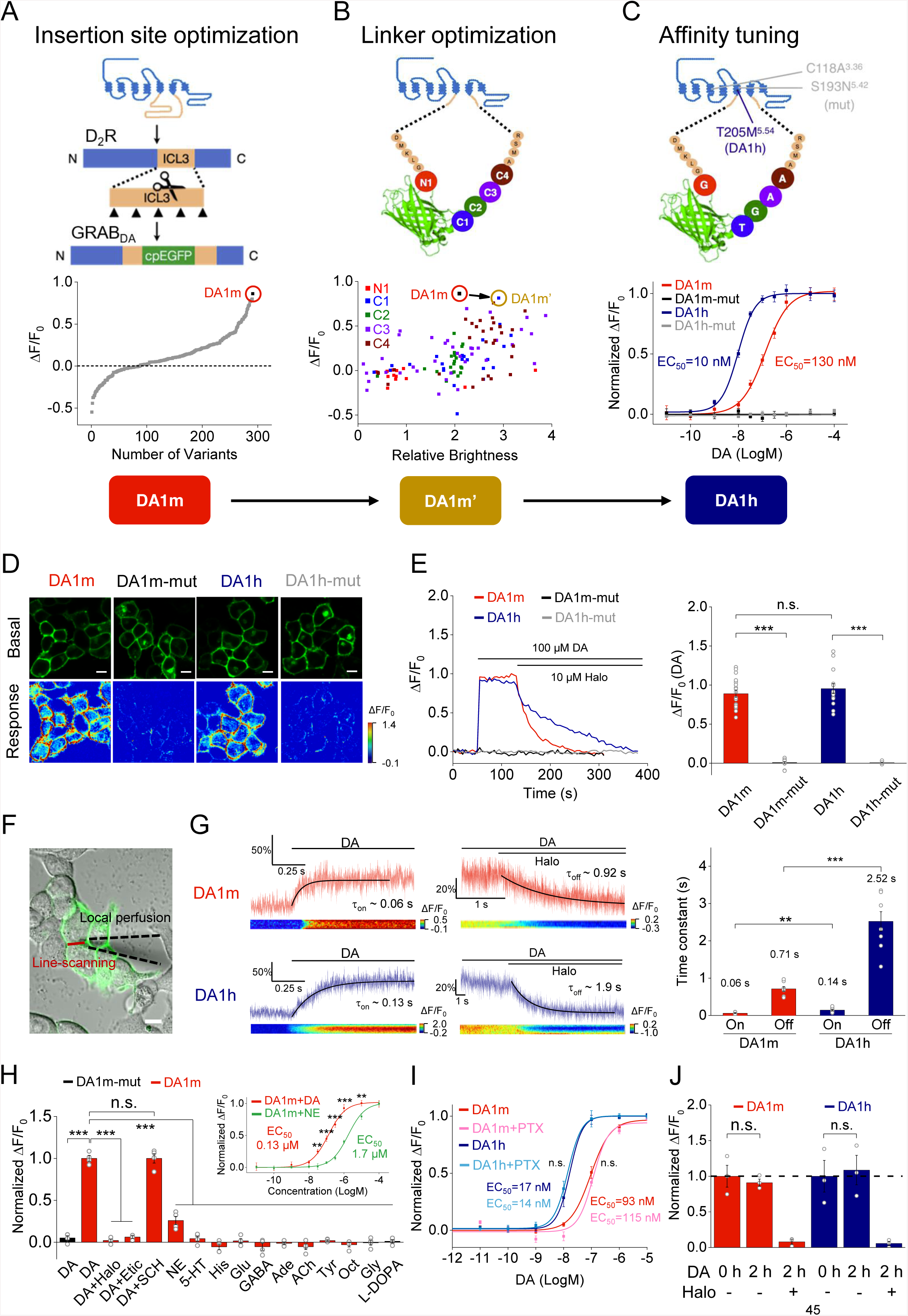
Design, optimization and characterization of GRAB_DA_ sensors in cultured HEK293T cells. (A-C) Schematic diagrams showing the strategy used to develop GRAB_DA_ sensors (top panels), and the corresponding performance of each variants in each optimization step (bottom panels). (A) Optimization of the cpEGFP insertion site within the third intracellular loop (ICL3) in D_2_R. The ΔF/F_0_ of GRAB_DA_-expressing cells in response to 100 μM DA application is shown below. GRAB_DA1m_, with the highest ΔF/F_0_ (∼90%), was selected for further optimization. Each data point represents the average of 3-5 cells. (B) Optimization of the linkers between the D_2_R and cpEGFP. Mutants were generated by changing each linker residue to 20 possible amino acids. The ΔF/F_0_ of the GRAB_DA_-expressing cells relative to the brightness in response to 100 μM DA application is shown below. Each data point represents the average of 100-400 cells. (C) Affinity tuning. Either the T205M single mutation, or the C118A/S193N double mutations, were introduced into GRAB_DA_, and the normalized dose-dependent fluorescence responses of various GRAB_DA_-expressing cells in response to DA application are plotted below. Each data point represents average of 6 wells containing 100-400 cells per well. (D-E) Fluorescence changes in GRAB_DA_-expressing cells in response to 100 μM DA followed by 10 μM Halo. Peak ΔF/F_0_ values in response to DA are summarized in the right panel in (E) (GRAB_DA1m_: *n =* 18 cells from 4 cultures (18/4); GRAB_DA1m-mut_: *n =* 15/3; GRAB_DA1h_: *n =* 14/3; GRAB_DA1h-mut_: *n =* 14/3; *p* < 0.001 between DA1m and DA1m-mut; *p* < 0.001 between DA1h and DA1h-mut; *p* = 0.42 between DA1m and DA1h). (F) Schematic image showing the local perfusion system. A glass pipette (black dashed lines) filled with DA or Halo was positioned close to the GRAB_DA_-expressing cells, and fluorescence was measured using confocal line-scanning (red line). (G) Left and middle: fluorescence changes in GRAB_DA_-expressing cells in response to the local perfusion (on rate: 100 μM DA in pipette with normal bath solution; off rate: 1 mM Halo in pipette with bath solution containing 10 μM DA for GRAB_DA1m_ or 1 μM DA for GRAB_DA1h_). The traces are the average of 3 different ROIs on the scanning line and are shaded with ± SEM. Right: group data summarizing the response kinetics of GRAB_DA_-expressing cells in response to DA (on) or Halo (off) (*n =* 8/group; *p* = 0.0093 between on kinetics; *p* < 0.001 between off kinetics). (H) Normalized fluorescence changes in GRAB_DA1m_-and GRAB_DA1m-mut_-expressing cells in response to the application of indicated compounds at 1 μM, including: DA, DA + Halo, DA + Etic, DA + SCH-23390 (SCH), norepinephrine (NE), 5-HT, histamine (His), glutamate (Glu), gamma-aminobutyric acid (GABA), adenosine (Ade), acetylcholine (ACh), tyramine (Tyr), octopamine (Oct), glycine (Gly), or L-DOPA (the first bar shows GRAB_DA1m-mut_-expressing cells in response to DA; *n* = 4 wells per group with 200-400 cells per well; *p* < 0.001 for DA-induced responses between GRAB_DA1m_ and GRAB_DA1m-mut_; *p* = 0.99 for GRAB_DA1m_ responses induced by DA comparing with DA+SCH; *p* < 0.001 for GRAB_DA1m_ responses induced by DA comparing with DA+Halo, DA+Etic, NE, 5-HT, His, Glu, GABA, Ade, ACh, Tyr, Oct, Gly and L-DOPA). The inset shows the normalized dose-dependent fluorescence responses of GRAB_DA1m_-expressing cells in response to DA (red) and NE (green) application (*n* = 6 wells per group with 100-300 cells per well; *p* = 0.002 at −7.5; *p* < 0.001 at −7, −6.5 and −6; *p* = 0.007 at −5). (I) Normalized fluorescence changes in GRAB_DA_-expressing cells in response to the application of DA, with or without the co-expression of pertussis toxin (PTX) (GRAB_DA1m_: *n* = 14/3; GRAB_DA1m_+PTX: *n* = 14/3; GRAB_DA1h_: *n* = 10/3; GRAB_DA1h_+PTX: *n* = 10/3; *p* = 0.680 comparing the EC_50_ of GRAB_DA1m_ and GRAB_DA1m_ +PTX; *p* = 0.810 for comparing the EC_50_ of GRAB_DA1h_ and GRAB_DA1h_ +PTX). (J) Normalized Fluorescence changes in GRAB_DA_-expressing cells during a 2-hour application of 100 μM DA (*n* = 3 wells/group; *p* = 0.620 for GRAB_DA1m_; *p* = 0.792 for GRAB_DA1h_). Scale bars, 10 μm in (D) and (F). Values with error bars indicate mean ± SEM. Students’ t-test performed; n.s., not significant; **, *p* < 0.01; ***, *p* < 0.001. See also Fig. S1-S3.

We next characterized the properties of the two GRAB_DA_ sensors in detail in cultured HEK293T cells. Both GRAB_DA1m_ and GRAB_DA1h_ trafficked efficiently to the plasma membrane of HEK293T cells (Fig. 1D and Fig. S1C). Cells expressing GRAB_DA1m_ and GRAB_DA1h_ exhibited robust fluorescence increases upon bath application of DA (Fig. 1D-F). Notably, the DA-induced fluorescence increase was completely blocked by the co-application of the D_2_R antagonist haloperidol (Halo) (Sokoloff et al., 1990), confirming the molecular specificity of these sensors (Fig. 1E). Both mutant forms of GRAB_DA_ sensors trafficked normally to the plasma membrane, similar to GRAB_DA1m_ and GRAB_DA1h_ (Fig. 1D and Fig. S1C), albeit with no detectable fluorescence increase in response to DA application (Fig. 1D,E). To characterize the response kinetics of GRAB_DA1m_ and GRAB_DA1h_, agonists or antagonists at high concentrations were locally applied to HEK293T cells expressing these sensors via a rapid perfusion system (Fig. 1F,G). Both GRAB_DA1m_ and GRAB_DA1h_ showed rapid fluorescence increases (on rate) in response to DA application, with a time constant of ∼100 ms (60 ± 10 ms for GRAB_DA1m_ and 140 ± 20 ms for GRAB_DA1h_, respectively), implying that GRAB_DA_ sensors are suitable for tracking rapid DA dynamics. The fluorescence decrease (off rate) of the GRAB_DA_ sensors in response to applications of the antagonist Halo is slower in GRAB_DA1h_ (2.5 ± 0.3 s) compared with GRAB_DA1m_ (0.7 ± 0.06 s), consistent with the differences in affinity (Fig. 1G). We also measured the photostability of GRAB_DA1m_ and GRAB_DA1h_ in HEK293T cells. Under confocal laser illumination, both GRAB_DA_ sensors were significantly more photostable than a membrane targeted EGFP or the glutamate sensor iGluSnFR (Marvin et al., 2013) (Fig. S2), suggesting that these new DA sensors are suitable for long-term measurements of DA dynamics.

We also tested the specificity of GRAB_DA_ sensors for DA comparing with other neurotransmitters. HEK293T cells expressing GRAB_DA_ sensors were sequentially perfused with solutions containing different neurotransmitters (Fig. 1H and Fig. S3; all were applied at 1 μM concentration). Again, bath application of DA induced robust fluorescence increases in both GRAB_DA1m_-and GRAB_DA1h_-expressing cells, and these signals were completely blocked by co-application of D_2_R antagonists Halo or eticlopride (Etic), but not by the D_1_R antagonist SCH-23390 (SCH). In contrast, applications of various other neurotransmitters, including serotonin (5-HT), histamine (His), glutamate (Glu), GABA, adenosine (Ade), acetylcholine (ACh), tyramine (Tyr), octopamine (Oct), glycine (Gly), or the DA precursor L-DOPA, did not elicit any detectable fluorescence changes (Fig. 1H and Fig. S3). We found that NE application evoked a small yet significant increase in fluorescence in both GRAB_DA1m_- and GRAB_DA1h_-expressing cells, although this increase was considerable smaller than the response elicited by DA applied at the same concentration (for GRAB_DA1m_, the ΔF/F_0_ in response to NE is 26% of that to DA). Further characterization of the affinity of GRAB_DA1m_ or GRAB_DA1h_ to both DA and NE revealed a 10-fold higher affinity to DA against NE (Fig. 1H and Fig. S3), as expected from the specificity of native human D_2_R to DA and NE (Lanau et al., 1997b). Overall, GRAB_DA1m_ and GRAB_DA1h_ sensors show rapid and highly sensitive responses to physiological ranges of DA with little or no sensitivity to almost all other neurotransmitters tested here.

Does ectopic expression of GRAB_DA_ sensors elicit unintended signaling by coupling to GPCR’s downstream pathways? To examine this, we separately tested the coupling efficacies of GRAB_DA_ sensors to either G protein-or β-arrestin-dependent pathways that are known to be downstream of activated D_2_R (Beaulieu and Gainetdinov, 2011). It is well established that the coupling of a G protein with its cognate GPCR significantly increases the receptor’s binding affinity for its ligand (Kruse et al., 2013). We therefore used pertussis toxin (PTX), which potently and selectively blocks the coupling between Gi proteins and GPCRs, such as D_2_R (Burns, 1988). In cells expressing either GRAB_DA1m_ or GRAB_DA1h_, co-expression of PTX did not significantly alter the sensors’ affinity for DA, indicating that GRAB_DA_ sensors do not couple extensively to Gi proteins (Fig. 1I). On the other hand, if the DA-binding of GRAB_DA_ sensors activates the β-arrestin-dependent pathway, the resulting internalization of sensors would reduce the fluorescence at the plasma membrane of HEK293T cells (Luttrell and Lefkowitz, 2002). However, we observed stable membrane fluorescence of the GRAB_DA_ sensor-expressing cells throughout a 2-hour exposure to DA at a saturation concentration (100 μM), suggesting no detectable activation of β-arrestin-dependent signaling under these conditions (Fig. 1J). Collectively, these data suggest that GRAB_DA_ sensors do not engage these major downstream GPCR-mediated signaling pathways.

### Characterization of GRAB_DA_ in cultured neurons

We next evaluated the expression pattern and functional properties of GRAB_DA_ sensors in cultured rat cortical neurons. After 48h expression, both GRAB_DA1m_ and GRAB_DA1h_ showed basal fluorescence signals in transfected neurons, with the brightest signal in the plasma membrane of somas and neurites, indicated by the colocalization with a membrane marker RFP-CAAX (Hancock et al., 1991) (Fig. 2A,B and Fig. S4). In addition, co-expression of the GRAB_DA_ sensors with the postsynaptic (PSD95-RFP) or presynaptic (SYP-RFP) markers revealed that GRAB_DA_ sensors were distributed throughout the entire membrane in dendritic shafts, spines, axons and axonal boutons, suggesting the suitability of GRAB_DA_ sensors to detect DA dynamics in sub-neuronal compartments (Fig. 2B and Fig. S4). As in HEK293T cells, DA application in transfected cultured neurons induced dose-dependent fluorescence increases in both GRAB_DA1m_-and GRAB_DA1h_-expressing neurons with a similar maximum ΔF/F_0_ of ∼90%, and with an EC_50_ of ∼170 nM and ∼8 nM for GRAB_DA1m_ and GRAB_DA1h_, respectively (Fig. 2C-E). Moreover, the specificity of both GRAB_DA_ sensors for DA was similar in cultured neurons compared to HEK 293T cells (Fig. 2F). Finally, no detectable decrease of surface fluorescence signals of GRAB_DA_ sensors-expressing cells was observed in response to a 2-h continuous application of DA at a saturation concentration (100 μM) (Fig. 2G), suggesting that GRAB_DA_ sensors were not internalized over this time frame. Collectively, these data demonstrate that GRAB_DA_ sensors can monitor DA signals with high sensitivity and specificity in cultured rat cortical neurons.

**Figure 2:**
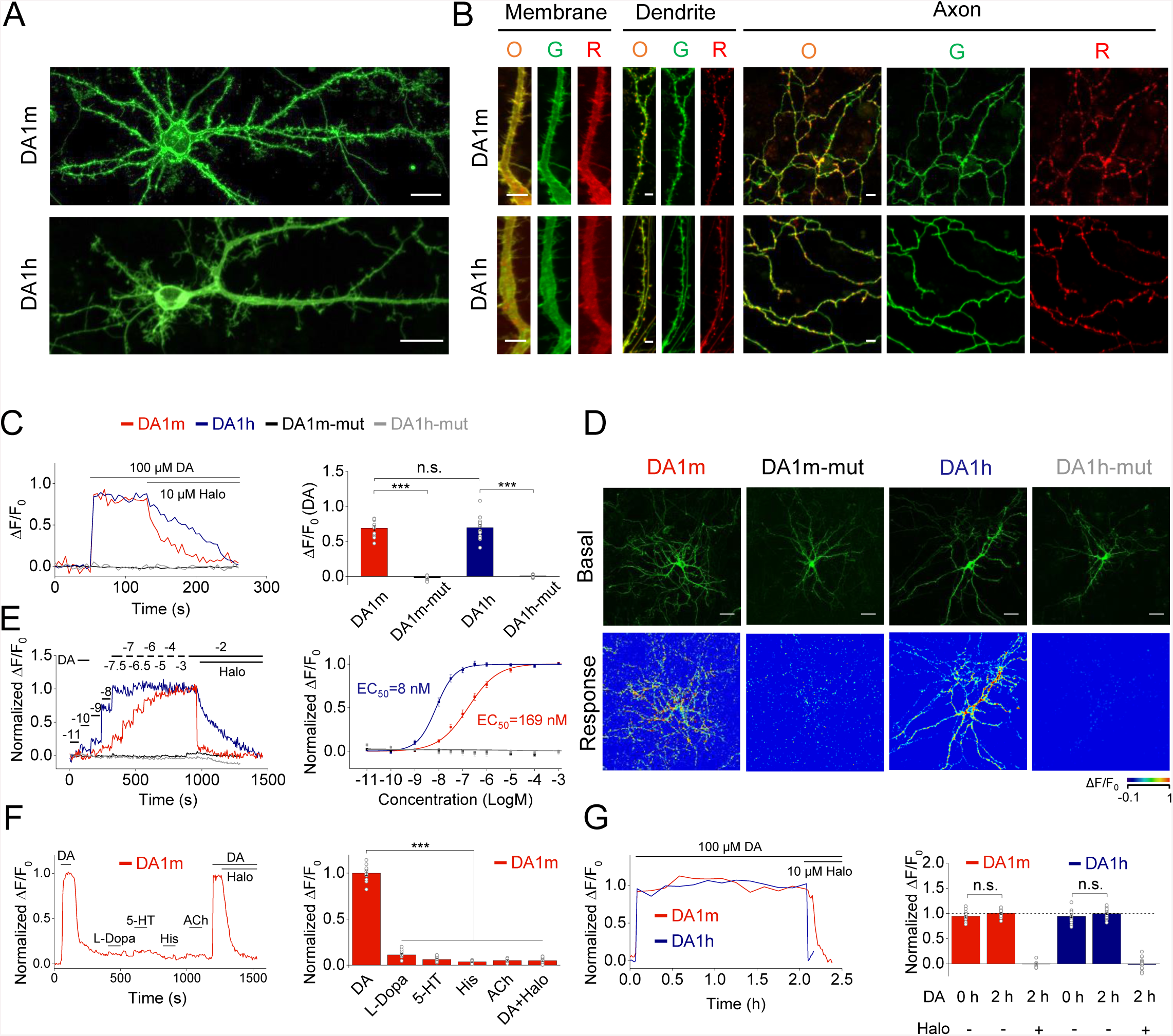
Characterization of GRAB_DA_ sensors in cultured neurons. (A) Expression of GRAB_DA_ sensors in cultured neurons. Scale bars, 20 μm. (B) Expression and localization of GRAB_DA_ sensors (green, G), subcellular markers (red, R) and overlay (O) in the indicated subcellular compartments in cultured neurons. RFP-CAAX, PSD95-RFP and Synaptophysin-RFP were co-expressed as markers of the plasma membrane, dendritic spines, and presynaptic boutons, respectively. Scale bars, 5 μm. (C and D) Fluorescence changes in GRAB_DA_-expressing neurons in response to the application of 100 μM DA followed by 10 μM Halo. Scale bars, 30 μm (GRAB_DA1m_: *n* = 13/7; GRAB_DA1m-mut_: *n* = 14/5; GRAB_DA1h_: *n* = 16/4; GRAB_DA1h-mut_: *n* = 10/5; *p* < 0.001 between DA1m and DA1m-mut; *p* < 0.001 between DA1h and DA1h-mut; *p* = 0.88 between DA1m and DA1h). (E) Time courses (left) and dose-dependent fluorescence changes (right) of GRAB_DA_-expressing neurons in response to DA application (GRAB_DA1m_: *n* = 10/6; GRAB_DA1m-mut_: *n* = 6/6; GRAB_DA1h_: *n* = 10/5; GRAB_DA1h-mut_: *n* = 10/3). (F) Fluorescence changes in GRAB_DA1m_-expressing neurons in response to the transient application of the indicated compounds at 1 μM, including DA, L-Dopa, 5-HT, His, ACh, DA(2^nd^) and DA+Halo (*n* = 12/12; *p* < 0.001 comparing responses in DA with that in L-Dopa, 5-HT, His, ACh and DA+Halo). (G) Fluorescence changes in GRAB_DA_-expressing neurons during a 2-hour application of 100 μM DA (GRAB_DA1m_: *n* = 20/12; GRAB_DA1h_: *n* = 14/6; *p* = 0.085 for DA1m; *p* = 0.085 for DA1h). Values with error bars indicate mean ± SEM. Student’s t-test performed; n.s., not significant; ***, *p* < 0.001. See also Fig. S4.

### Characterization of GRAB_DA_ in acute brain slices

A crucial issue is whether GRAB_DA_ sensors are sufficiently sensitive to detect and monitor endogenous DA release in native brain tissue. To begin to resolve this issue, we injected AAVs carrying either GRAB_DA1m_ or GRAB_DA1h_ into the nucleus accumbens (NAc) of mice, then prepared acute brain slices containing NAc two weeks later. Post hoc visualization in fixed brain slices revealed that the fluorescent signal of GRAB_DA_ sensors could be detected in the NAc region of virus-injected mice, and the signal was in close proximity to putative DA-releasing fibers that were immunoreactive for tyrosine hydroxylase (TH) (Fig. 3A). Electrical stimulation within NAc core elicited robust and time-locked fluorescence increases in both GRAB_DA1m_-and GRAB_DA1h_-expressing neurons (Fig. 3A-C). Increasing the number of stimulation pulses or stimulation frequency resulted in a progressive increase in the intensity of evoked fluorescence signals in neurons expressing GRAB_DA1m_ or GRAB_DA1h_ (Fig. 3B,C). These stimulus-evoked fluorescent signals reached plateau levels at high stimulation frequency with large stimulation number (e.g. more than 50 pulses) in GRAB_DA1h_ expressing neurons, presumably due to saturation of the sensor arising from GRAB_DA1h_’s high affinity for DA. The rise times of stimulus-evoked fluorescence signals were fast in both GRAB_DA1m_-and GRAB_DA1h_-expressing neurons (Fig. 3D), whereas the decay time of fluorescent signals in GRAB_DA1h_ expressing neurons was slower than that in neurons expressing GRAB_DA1m_ (Fig. 3D), consistent with the DA affinity and response kinetics measured in cultured cells (see Fig. 1C,G). Repeated electrical stimuli delivered at 5-minute intervals evoked reproducible fluorescence responses, indicating the reliability and stability of GRAB_DA1m_ and GRAB_DA1h_ in reporting multiple DA releasing events (Fig. 3E). Bath application of the D_2_R antagonist Halo abolished the electrically stimuli-induced fluorescence responses in either GRAB_DA1m_-or GRAB_DA1h_-expressing neurons (Fig. 3F), verifying the sensor’s specificity when expressed in brain slices. Collectively, both GRAB_DA1m_ and GRAB_DA1h_ enable sensitive and specific detection of endogenous DA dynamics in acute mouse brain slices.

**Figure 3:**
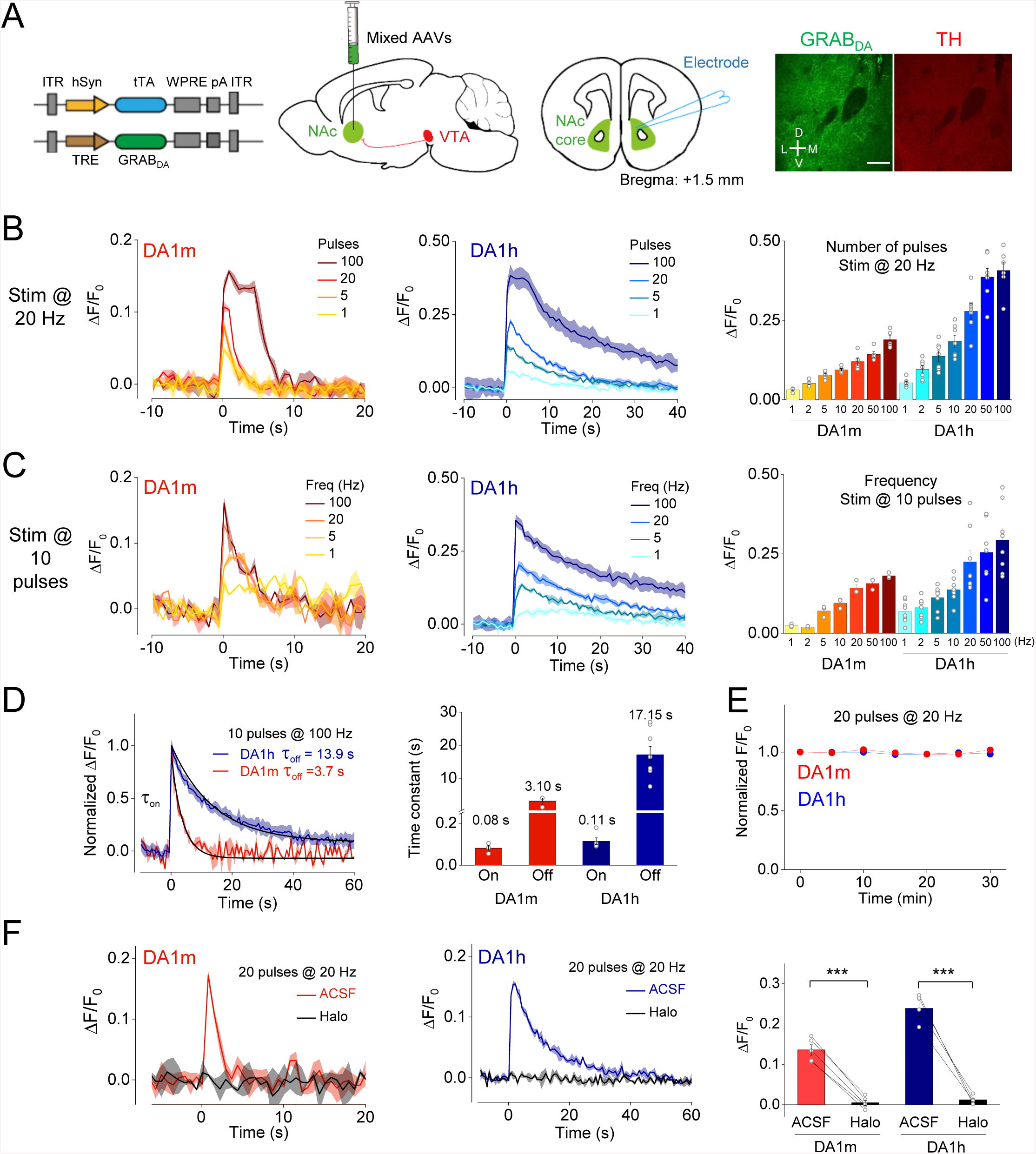
Release of endogenous DA measured in acute mouse brain slices. (A) Left three panels, schematic diagrams of the viral expression vector, experimental protocol for expressing GRAB_DA_ sensors and imaging DA dynamics in mouse brain slices containing NAc. Right, the immunoreactive signals of GRAB_DA_ (green) expressed in NAc neurons and TH (red) in dopaminergic terminals. Scale bar, 100 μm. (B) Representative traces (left and middle) and group analysis (right) of the fluorescence changes in GRAB_DA1m_-and GRAB_DA1h_-expressing neurons in response to a train of 20 Hz electrical stimuli containing the indicated pulse numbers. Each trace is the average of 3 separate trials in one slice (GRAB_DA1m_: *n* = 5 slices from 3 mice; GRAB_DA1h_: *n* = 7 slices from 4 mice). (C) Similar as (B), except that a train of 10-pulse electrical stimuli was applied at the indicated frequencies (GRAB_DA1m_: *n* = 3 slices from 2 mice; GRAB_DA1h_: *n* = 8 slices from 4 mice). (D) Representative traces (left) and group analysis (right) of the normalized fluorescence changes and kinetics in GRAB_DA1m_-and GRAB_DA1h_-expressing neurons in response to 10 electrical pulses delivered at 100 Hz. The rising (on) and decaying (off) phases in the traces were fitted separately, and the response time constants are summarized on the right (GRAB_DA1m_: *n* = 3 slices from 2 mice; GRAB_DA1h_: *n* = 5-8 slices from 3 mice). (E) The fluorescence changes in GRAB_DA1m_-and GRAB_DA1h_-expressing neurons in response to multiple trains of electrical stimuli at an interval of 5 min. The fluorescence changes measured during the first train in each slice were used to normalize the data (GRAB_DA1m_: *n* = 3 slices form 2 mice; GRAB_DA1h_: *n* = 6 slices from 3 mice). (F) Representative traces (left and middle) and group analysis (right) of the fluorescence changes in GRAB_DA1m_-and GRAB_DA1h_-expressing neurons in response to 20 electrical pulses (at 20 Hz), in control solution (ACSF) or solution containing 10 μM Halo (GRAB_DA1m_: *n* = 5 slices from 4 mice, *p* < 0.001 comparing ACSF with Halo; GRAB_DA1h_: *n* = 6 slices from 4 mice, *p* < 0.001 comparing ACSF with Halo). Values with error bars indicate mean ± SEM. The shaded areas and error bars indicate ± SEM. Student’s t-test performed; ***, *p* < 0.001.

### Imaging DA dynamics in *Drosophila*

We next examined the ability of GRAB_DA_ sensors to detect physiologically relevant DA dynamics in living animals. We selected the fly as an initial test of our sensors, as DA plays a key role in the fly brain, serving as a critical teaching signal in olfactory-associative learning (Burke et al., 2012; Heisenberg, 2003; Liu et al., 2012; Schwaerzel et al., 2003). Transgenic UAS-GRAB_DA1m_ flies were generated and first crossed with TH-GAL4 to express GRAB_DA1m_ specifically in DANs. Two-photon imaging methods were used to study odor-evoked DA signals in the mushroom body (MB) in living flies (Fig. 4A). We found that the odorant isoamyl acetate (IA) elicited a time-locked fluorescence increase in the MB, most prominently in the β’ lobes, and this IA-evoked response was blocked by Halo application. In contrast, no IA-evoked fluorescence response was observed in flies expressing GRAB_DA1m-mut_ (Fig. 4B). To further test the signal specificity, we use the C305a-GAL4 line to express GRAB_DA1m_ in Kenyon cells which receives direct input from DANs (Aso et al., 2010). Comparing wild-type (WT) control flies and TH-deficient flies (Cichewicz et al., 2017), which lack DA synthesis in the CNS. Compared with WT flies, we observed no detectable IA-evoked GRABDA1m fluorescence increase was observed in TH-deficient flies (Fig. 4B,C). We also examined whether the ectopic expression of GRABDA sensors may alter physiological properties, such as the neuronal excitability. We observed no significant difference between GRABDA sensors-expressing and non-expressing DANs or Kenyon cells with respect to odor-evoked Ca^2+^ signals (Fig. S5), suggesting that expression of the sensor does not alter odor-evoked responses in neurons in the fly brain.

**Figure 4:**
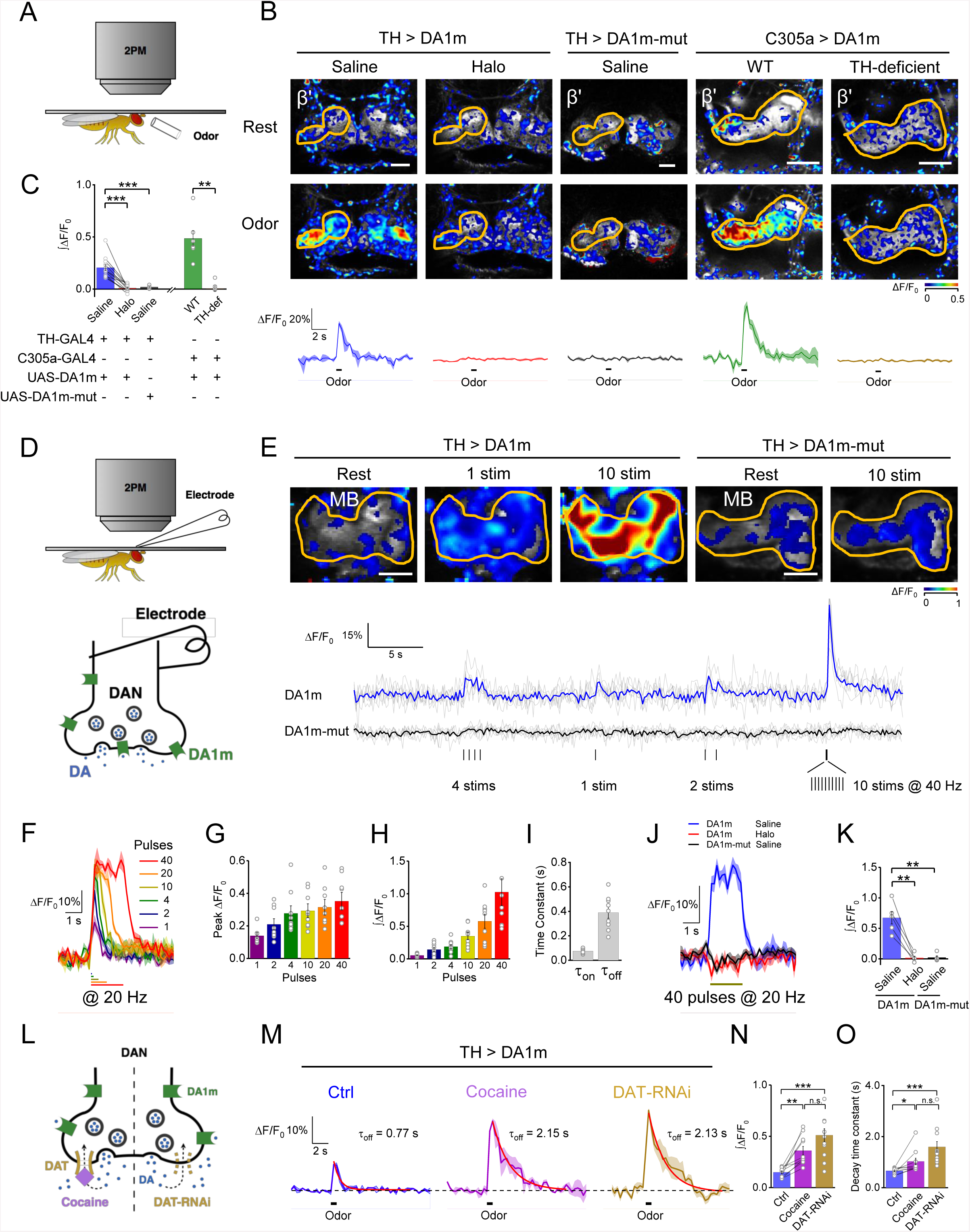
*In vivo* imaging of DA dynamics in the *Drosophila* brain. (A) Schematic illustration depicting the *in vivo* olfactory stimulation and imaging experiment under two-photon microscopy. (B and C) Representative pseudo-color images and traces (B) and group analysis (C) of the fluorescence changes of TH > GRAB_DA1m_ and TH > GRAB_DA1m-mut_ flies in response to 1 s olfactory stimulation (TH > GRAB_DA1m_: *n =* 12 flies; TH > GRAB_DA1m-mut_: *n =* 5 flies; c305a > GRAB_DA1m_ with WT background: *n =* 6 flies; c305a > GRAB_DA1m_ with TH-deficient background: *n =* 6 flies; *p* < 0.001 for responses of TH > GRAB_DA1m_ in saline comparing with Halo; *p* < 0.001 for responses of TH > GRAB_DA1m_ in saline comparing with TH > GRAB_DA1m-mut_ in saline; *p* = 0.002 for responses of c305a > GRAB_DA1m_ with WT background comparing with TH-deficient background). (D) Schematic illustrations depicting *in vivo* electrical stimulation experiment, in which the electrode was positioned near the GRAB_DA1m_-expressing DANs in order to evoke DA release. (E) Representative pseudo-color images of the fluorescence changes in TH > GRAB_DA1m_ and TH > GRAB_DA1m-mut_ flies in response to multiple trains of electrical stimuli. Shown below are single-trial traces (in gray) and 6-trial averaged traces (blue and black) measured in one fly with indicated genotypes. Each vertical tick indicates 1 ms electrical stimulation (“stim”). (F-I) Fluorescence changes in TH > GRAB_DA1m_ flies in response to the indicated pulses of electrical stimuli (at 20 Hz), showing representative traces (F), group data for peak ΔF/F_0_ (G), integrated data (H) and the response kinetics (I) (*n* = 9 flies per group). (J and K) Representative traces (J) and group analysis (K) of fluorescence changes in TH > GRAB_DA1m_ and TH > GRAB_DA1m-mut_ flies in response to 40 pulses electrical stimuli (at 20 Hz), in normal saline or in saline containing 10 μM Halo (TH > GRAB_DA1m_: *n =* 5 flies; TH > GRAB_DA1m-mut_: *n =* 5 flies; *p* = 0.004 for responses of TH > GRAB_DA1m_ in saline comparing with Halo; *p* = 0.007 for responses of TH > GRAB_DA1m_ in saline comparing with TH > GRAB_DA1m-mut_ in saline). (L-O) Fluorescence changes in TH > GRAB_DA1m_ flies in response to 1 s olfactory stimulation, in control condition, in the presence of the DAT blocker cocaine (3 μM) or when the DAT expression in DAN was impaired by DAT-RNAi. Schematic illustration of the experimental design in (L). Representative traces (M) were fitted with a single-exponential function (red traces), with the decay time constants shown. The group analysis of integrated data and the decay time constants are shown in (N) and (O), respectively (TH > GRAB_DA1m_: *n =* 10 flies; TH > GRAB_DA1m_, DAT-RNAi: *n =* 11 flies; between control and cocaine groups, *p* = 0.002 for integrals and *p =* 0.025 for decay time constants; between control and DAT-RNAi groups, *p* < 0.001 for both integrals and decay time constants; between cocaine and DAT-RNAi groups, *p* = 0.095 for integrals and *p* = 0.053 for decay time constants). The fluorescence traces in (B), (F), (J), and (M) are the averaged results of 3∼6 trials from one fly, and the shaded area indicates ± SEM. Values with error bars indicate mean ± SEM. Student’s t-test performed; n.s., not significant; *, *p* < 0.05; **, *p* < 0.01; ***, *p* < 0.001. Scale bars in (B) and (E), 25 μm. See also Fig. S5.

To further characterize the sensitivity and kinetics of GRAB_DA1m_ *in vivo*, we electrically stimulated MB DANs while simultaneously monitoring the corresponding fluorescence signals (Fig. 4D,E). We found that GRAB_DA1m_-expressing DANs exhibited a reproducible fluorescence increases in response to repeated trains of electrical stimuli delivered at various frequencies, and even a single stimulus was sufficient to elicit a measurable increase in fluorescence (Fig. 4D,E). In contrast, similar electrical stimulation did not evoke any detectable fluorescence increase in flies expressing GRAB_DA1m-mut_ (Fig. 4D,E). With the increase of pulse number delivered at 20 Hz, fluorescence signals of GRAB_DA1m_ increased progressively and reached a plateau of peak ΔF/F_0_ of ∼35% after 10 pulses (Fig. 4F-H). The fluorescence responses evoked by electrical stimulation exhibited sub-second kinetics, with on and off time constants of 0.07 ± 0.01 s and 0.39 ± 0.05 s, respectively (Fig. 4I), indicating the suitability of GRAB_DA1m_ for monitoring endogenous transient DA signals. Consistent with the odor-evoked fluorescence increase, application of Halo completely blocked the fluorescence increases elicited by electrical stimuli (Fig. 4J,K), confirming the sensor’s specificity.

The DA transporter (DAT), which is a highly conserved protein across insects and vertebrates, is responsible for the reuptake of DA from extracellular space for subsequent reuse, and is the primary target of many drugs of abuse (Bainton et al., 2000; Ritz et al., 1987). Impairing DAT function results in sustained elevation of extracellular DA levels (Fig. 4L) (Bainton et al., 2000; Ritz et al., 1987). To determine whether our sensor could detect changes in extracellular DA arising from manipulation of the DAT, we applied cocaine, a psychostimulant drug that blocks DAT. Cocaine significantly potentiated the odor-evoked fluorescence increase of GRAB_DA1m_ expressed in MB DANs, and this cocaine-potentiated response was accompanied by a prolonged decay in the fluorescence signal (Fig. 4M-O). Genetically knocking down the expression of DAT selectively in DANs phenocopied the effect of cocaine administration (Fig. 4M-O), confirming the ability of GRAB_DA1m_ to measure the dynamic regulation of DA release and reuptake *in vivo*. Together with our results described above, these data demonstrated that GRABDA sensors have the sensitivity, fast kinetics, and specificity to report *in vivo* DA dynamics in genetically-defined neurons in the intact brain of living flies.

### Imaging DA release in the intact zebrafish brain

Zebrafish larvae have an optically transparent brain and is capable of performing a wide range of behaviors, affording a powerful system to explore the structure and function of the vertebrate brain at cellular resolution in a behavioral framework. To test the feasibility of using GRAB_DA_ sensors in imaging DA dynamics in zebrafish larvae, we generated the transgenic line Tg(elval3:GRAB_DA1m_/DAT:TRPV1-TagRFP), in which GRAB_DA1m_ was expressed pan-neuronally throughout the brain, and TRPV1-TagRFP was expressed specifically in DANs to enable their chemogenetic activation by capsaicin (Fig. 5A). We first applied DA to the fish, and observed fluorescence increases in GRAB_DA1m_-expressing neurons in the head, which were blocked by co-application of antagonist Halo, suggesting the specificity of GRAB_DA_ (Fig. 5B-D).

**Figure 5:**
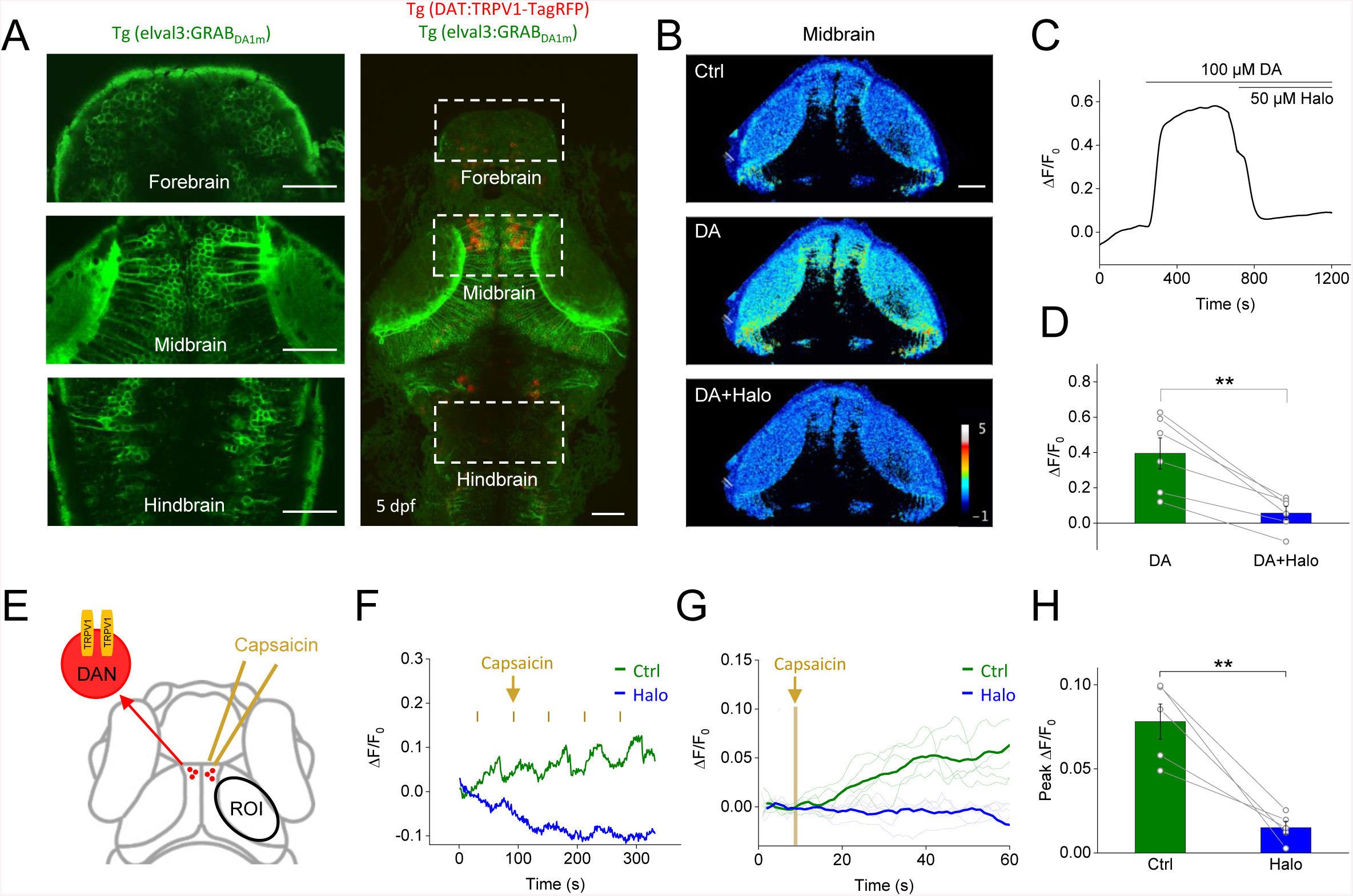
Monitoring *in vivo* DA release in transgenic zebrafish. (A) Fluorescence images of a transgenic zebrafish larvae expressing GRAB_DA1m_ (green) pan-neuronally and TRPV1-TagRFP (red) in DANs. Zoom-in view of GRAB_DA1m_-expressing neurons in indicated brain regions are shown (left). (B-D) Representative pseudo-color images (B), trace (C) and group analysis (D) of fluorescence changes of GRAB_DA1m_-expressing neurons in response to 100 μM DA followed by 50 μM Halo (*n* = 6 fishes; *p* = 0.002 between DA and DA+Halo). (E) Schematic diagram showing the experimental design to chemogenetically activate TRPV1-expressing DANs by capsaicin. The tectal neurons downstream of the DANs were analyzed as indicated by the region of interest (ROI). (F) Representative traces of the fluorescence changes in GRAB_DA1m_-expressing neurons in response to 5 pulses of 50 μM capsaicin (each vertical orange tick indicates 100-ms puffs of capsaicin with a 1 min interval) in control normal solution (green) or solution containing 50 μM Halo (blue). (G and H) Averaged traces (G) and group analysis (H) of the fluorescence changes in GRAB_DA1m_-expressing neurons in response to single trial of capsaicin application, in control normal solution (green) or solution containing 50 μM Halo (blue) (*n =* 5 fishes; *p* = 0.006 between control and Halo). Scale bars in (A) and (B), 50 μm. Paired student’s t-test performed.

Next, we hope to track the dynamics of endogenous DA signals in zebrafish larvae. We previously reported that the optic tectum of zebrafish larvae receives synaptic input from pretectal DANs (Shang et al., 2015), advancing the tectum as a potential site to monitor DA release. We therefore performed confocal imaging of GRAB_DA1m_-expressing tectal neurons in the live transgenic zebrafish larvae (Fig. 5E). Repeated application of capsaicin (five 100-ms puffs delivered with a 1-min interval) caused the progressive increase in the fluorescence signals in the tectal neuropil (Fig. 5F-H). The capsaicin-induced fluorescence increase was again abolished by the application of Halo (Fig. 5F-H), confirming that the response was specifically due to GRAB_DA1m_ activation. Thus, the GRAB_DA1m_ is well suited to report *in vivo* DA dynamics in the brain of zebrafish larvae.

### Combining optogenetics with GRAB_DA_ to measure the dynamics of DA in freely moving mice

To test the ability of GRAB_DA_ sensors to report DA dynamics in specific circuits in the mouse brain *in vivo*, we focused on DANs located in the substantia nigra pars compacta (SNc) that project to the dorsal striatum (Str). This nigrostriatal pathway is implicated in complex behavioural functions including motivation, reward, and learning (Balleine et al., 2007; Da Silva et al., 2018; Howe and Dombeck, 2016). We virally expressed DIO-C1V1 (Yizhar et al., 2011) in the SNc of TH-Cre mice, permitting selective optogenetic activation of DANs. Additionally, we virally co-expressed GRAB_DA1m_ and tdTomato in the dorsal striatum, allowing us to simultaneously monitor DA release using the green (GRAB_DA1m_) channel while detecting movement-related fluorescence artefacts using the red (tdTomato) channel (Fig. 6A,B, see methods for detail). In free moving mice, the ratio of GRAB_DA1m_ to tdTomato fluorescence was elevated upon the administration of methylphenidate, a known DAT blocker (Volkow et al., 1999) (Fig. 6C, top), and was suppressed by subsequent administration of Etic (Fig. 6C, top), a D_2_R blocker, implying the ability of GRAB_DA_ sensors in reporting DA dynamics to pharmacological treatments. Interestingly, we observed fluctuations in the ratio of GRAB_DA1m_ to tdTomato fluorescence that likely reflect the spontaneous DA release during the animal movements (Balleine et al., 2007; Da Silva et al., 2018; Howe and Dombeck, 2016), as they were prolonged during methylphenidate application and largely diminished by Etic administration (Fig. 6C, bottom).

**Figure 6:**
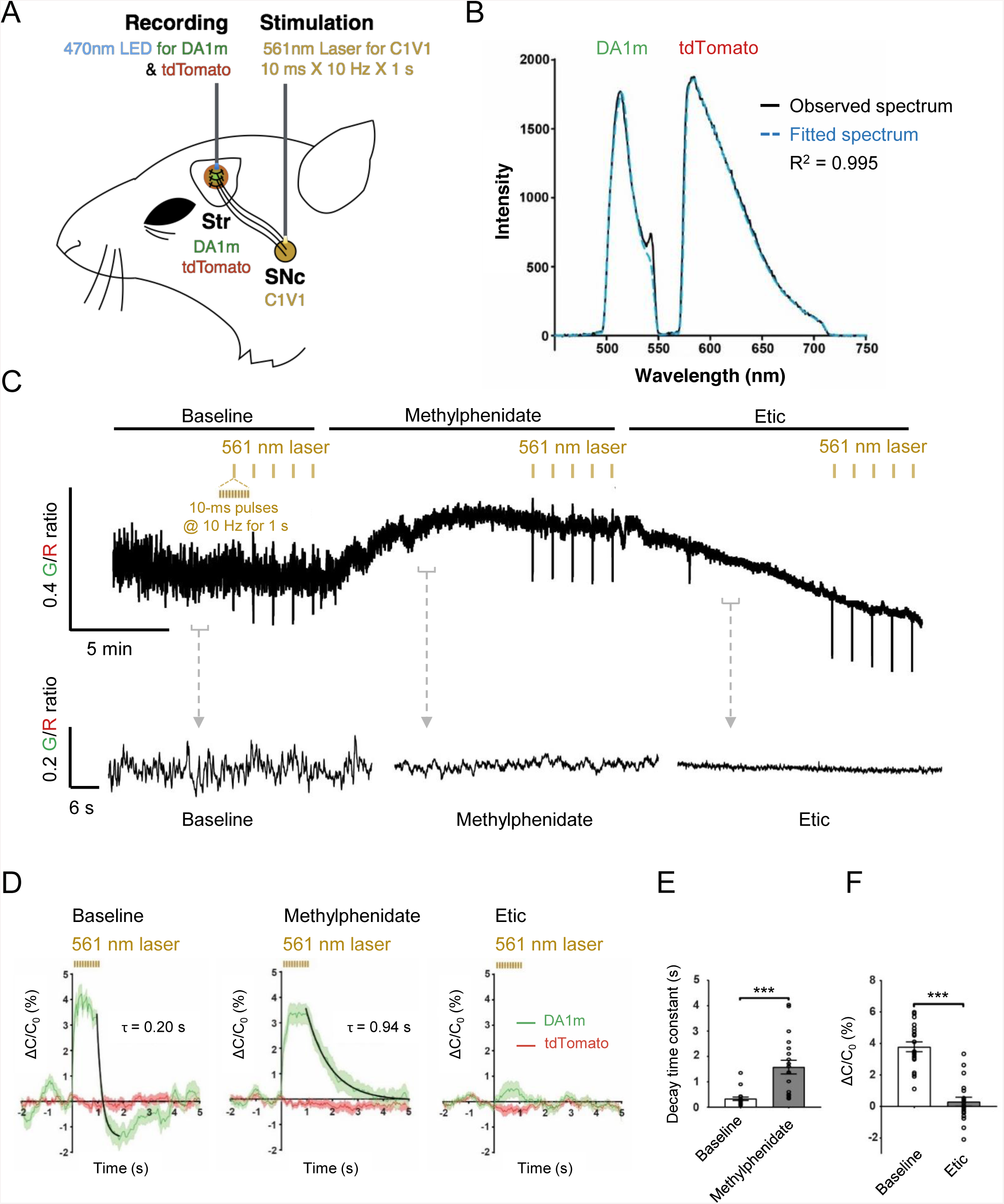
Striatal DA dynamics measured in freely moving mice during optogenetic stimulation of the SNc. (A) Illustration depicting the experimental design with dual-color optical recordings of GRAB_DA1m_-and tdTomato-expressing neurons in the dorsal striatum during simultaneous optogenetic stimulation of DANs in the SNc. (B) A representative frame of the emission spectra of GRAB_DA1m_ and tdTomato co-expressed in the dorsal striatum. The black trace shows the measured spectrum; the blue dashed trace shows the corresponding best fitting curve generated by a linear unmixing algorithm. (C) Spontaneous DA fluctuations represented by the ratio of GRAB_DA1m_ to tdTomato fluorescence in a freely moving mouse (top) and its representative traces in the control condition (bottom left), 5 min after the i.p. injection of DAT blocker methylphenidate (10 mg/kg, bottom middle), and 5 min after the i.p. injection of D_2_R antagonist Etic (2 mg/kg, bottom right). Black lines above indicate the time of compound administration. Yellow ticks indicate the time of optogenetic stimulation. (D) Averaged fluorescence changes (mean ± SEM, *n* = 20 trials from 4 hemispheres of 2 mice) of GRAB_DA1m_ (green) in the dorsal striatum in response to optogenetic C1V1 stimulation of DANs in the SNc under indicated conditions: baseline (left), after the i.p. injection of methylphenidate (middle), and after the i.p. injection of Etic (right). The off kinetics were fitted with a single-exponential function (black traces). (E) Comparison of the decay time constants of C1V1-evoked GRAB_DA1m_ fluorescence responses between the control condition and after methylphenidate injection (*n* = 20 trials from 4 hemispheres of 2 mice). (F) Comparison of the magnitude of C1V1-evoked GRAB_DA1m_ fluorescence changes between the control condition and after Etic injection (*n* = 20 trials from 4 hemispheres of 2 mice). ***, *p* < 0.001, student’s t-test performed in (E) and (F).

Combining optogenetic stimulation and fiber photometry in freely moving mice, we observed that C1V1-evoked activation of DANs in the SNc generated a time-locked transient fluorescence increase in the dorsal striatum specific to the GRAB_DA1m_ channel (Fig. 6D), consistent with the evoked DA release from SNc DAN terminals. Systemic administration of the DA transporter blocker methylphenidate significantly prolonged the decay of optogenetically-evoked GRAB_DA1m_ fluorescence responses in the dorsal striatum (Fig. 6D,E). Furthermore, administration of D_2_R antagonist Etic largely abolished the response (Fig. 6D,F). Therefore, the GRAB_DA_-and C1V1-based all-optical approach is effective for monitoring spontaneous and optogenetically-evoked DA release in the nigrostriatal pathway of mice.

### Bi-directional modulation of DA dynamics in the nucleus accumbens (NAc) during Pavlovian conditioning

In addition to the nigrostriatal pathway, the dopaminergic projection from the ventral tegmental area (VTA) to the nucleus accumbens (NAc) regulates a variety of important functions including the reinforcement learning (Daw and Tobler, 2013; Glimcher, 2011; Gonzales et al., 2004), wherein an animal learns to associate an initially neutral sensory cue, such as a short tone burst, with an ensuing reward or punishment. Previous studies show that DANs in the VTA fire phasically to unpredicted rewards or reward-predicting cues and, with conditioning, the reward-evoked response diminishes as it becomes fully predicted by the cue (Bayer and Glimcher, 2005; Schultz, 2006; Schultz et al., 1997). To test whether our sensor can detect behaviourally relevant changes in endogenous DA release, we first expressed GRAB_DA1h_ in the NAc of head-fixed, water-restricted mice and trained them to associate a brief auditory cue with subsequent delivery of a liquid sucrose water reward (Fig. 7A). Each mouse experienced two distinct auditory cues, one that predicted delivery of a reward within a variable delay of 500-1500 ms after the end of the cue, as well as a second, randomly interleaved control cue (No Water, N.W.) that was not associated with water reward. Mice were trained daily for ∼10 days, and the GRAB_DA1h_ signal was recorded using *in vivo* fiber photometry in both the early and late stages of training. Consistent with the expected DA signaling in this classical paradigm, elevations in the GRAB_DA1h_ fluorescent signal aligned to reward delivery in every mouse, immediately in the first session without requiring training (Fig. 7C,E). After 6-10 days of training, mice selectively learned to associate the reward-predictive cue with delivery of reward. Consistent with the established reward prediction error theory of DA function, DA signals emerged in trained mice in response to the reward-predicting cue before the delivery of the reward (Fig. 7B,D-F). These results demonstrate that the signal-to-noise and temporal resolution of GRAB_DA1h_ is sufficient to detect physiologically relevant DA signaling *in vivo* in awake, behaving mice. Furthermore, these results serve to validate the GRAB_DA1h_ recorded DA signals by anchoring them to decades of established basal ganglia and the physiology of DA neurons.

**Figure 7:**
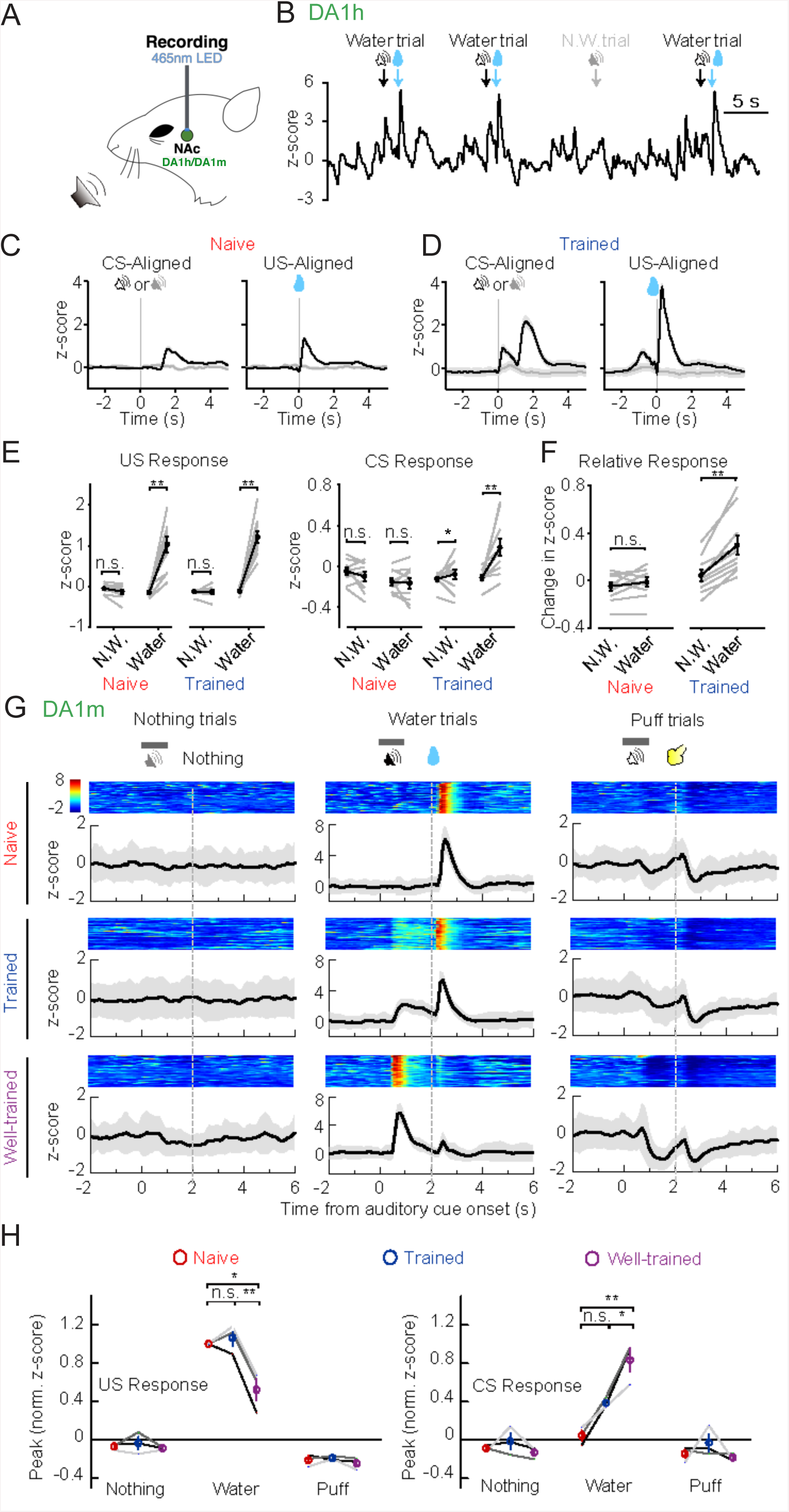
Dopamine release in NAc measured during various training phases of an auditory conditioning task. (A) Recording configuration for head-fixed Pavlovian conditioning task. Dopamine dynamics in the NAc were monitored by recording the fluorescence changes in GRAB_DA_-expressing neurons using fiber photometry. (B) Exemplar trace from *in vivo* fiber photometry recording from a trained mouse expressing GRAB_DA1h_ in NAc, encompassing four sequential trials. The timings of cues (CS) or water reward (US) are indicated above. (C) Task-aligned photometry signals from an exemplar mouse in the first behavioral session aligned to the cue (CS, left) or to the reward delivery (US, right). Note robust reward-related signal and absence of cue-related signal. (D) Task-aligned photometry signals from the same moues in (C) after training. Note emergence of DA response to reward-predictive cue. (E) Group data demonstrating GRAB_DA1h_ responses to water (US, left) or cue (CS, middle) in the NAc of both naïve and trained mice (*n* = 9 mice; US response: naïve N.W.: *p* = 0.084; naïve water: *p* = 0.0020; trained N.W.: *p* = 0.56; trained water: *p* = 0.0020; CS response: naïve N.W.: *p* = 0.37; naïve water: *p* = 1.0000; trained N.W.: *p* = 0.043; trained water: *p* = 0.0020, signed rank test performed). (F) Direct comparison of baseline-subtracted cue responses reveals equivalent lack of response to N.W. or reward-predictive cue in naïve mice (left), and elevated response to reward-predictive cue over N.W. cue in trained mice (right). Baseline subtraction was performed trial-by-trial by comparing the cue response (100 ms – 1000 ms following onset of the cue) to 900 ms before onset of the cue. This cue response window encompassed the auditory cue as well as the variable delay period and terminated before the earliest reward delivery times (naïve: *p* = 0.43; trained: *p* = 0.0020, signed rank test performed). (G) Representative fluorescence changes in GRAB_DA1m_-expressing neurons in the NAc during training. The heat map shows the fluorescence changes in the first 50 consecutive trials in each category, with each trial time-aligned to the onset of the auditory cue. The lower panels show the post-event histograms (PSTH) from all trials; the shaded area indicates ± SD. The fluorescence changes measured during naïve, trained and well-trained phases in the same mouse are shown, respectively. (H) Group analysis of the normalized peak z-scores for the US and CS in the indicated training phases. Each trace (coded with specific gray value) represents data from one animal (*n =* 3 mice; US: *p* = 0.7638 between naive and trained, *p* = 0.0125 between naive and well-trained, *p* = 0.0080 between trained and well-trained; CS: *p* = 0.1032 between naive and trained, *p* = 0.0067 between naive and well-trained, *p* = 0.0471 between trained and well-trained). Error bars, ± SEM. Post-hoc Tukey’s test was performed; n.s., not significant; *, *p* < 0.05; **, *p* < 0.01; ***, *p* < 0.001.

In addition to transient elevations in DA signal in response to unexpected reward, DA neuron firing dips briefly in response to aversive stimuli (Brooks and Berns, 2013; Schultz, 2007; Ungless et al., 2004). We expressed the lower affinity GRAB_DA1m_ in the NAc of mice and trained them to associate a tone burst with an ensuing reward (a drop of water), or a punishment (a brief air puff to the face) to test whether our sensor can detect bi-directional changes in DA tone. During training, mice learned the association between an auditory cue and several outcomes (Fig. 7A). *In vivo* fiber photometry recording of GRAB_DA_ signals revealed that during early stages of training, reward or punishment delivery triggered a robust increase or decrease in the fluorescence signal in the NAc (Fig. 7G,H). Over the course of training, the magnitude of this reward-evoked response decreased, while a response of similar sign developed to the associated cue (Fig. 7G,H). In summary, the GRAB_DA_ sensor can be used to report the dynamic bi-directional changes in DA release over the course of Pavlovian conditioning.

### Monitoring DA release in the NAc of mice during male mating behaviors

In contrast to the well-established involvement of DA in Pavlovian conditioning, DA dynamics during naturally rewarding social behaviors (Berridge and Robinson, 1998), such as courtship and mating, remains largely a subject of debate. Microdialysis and voltammetric measurements have previously revealed a relatively slow change in DA concentration during sexual behaviors: DA levels in the male’s NAc start to increase after a female is introduced into his cage and continues to rise throughout the course of sexual behaviors and ejaculation (∼20 minutes), then slowly (∼30 minutes) return to baseline after the female is removed (Damsma et al., 1992; Mas et al., 1995; Pfaus et al., 1990b). In contrast, a more recent study using FSCV showed that DA is transiently released in the male’s NAc when the female is introduced but few changes in DA levels were detected during subsequent sexual behaviors (Robinson et al., 2002; Robinson et al., 2001). To better understand DA dynamics during sexual behaviors, we virally expressed GRAB_DA1h_ in the NAc of male mice and used fiber photometry to record DA signals during sexual behaviors (Fig. 8A). Four weeks after injection, a sexually receptive C57BL/6 female was introduced into the home cage of the male mouse. Upon a female introduction, the male quickly approached and investigated the female, and then initiated mounting within the first minute. The GRAB_DA1h_ signals measured in the male’s NAc acutely and consistently increased with investigation of the female, mounting, intromission, ejaculation, and penile grooming (Fig. 8B). When we aligned the GRAB_DA1h_ signals with the manually annotated behaviors for each male, we observed that fluorescence signals increased prior to the corresponding behavior, peaked at the behavior’s onset, and then gradually declined (Fig. 8C). Among all of the sexual behaviors we annotated, the largest fluorescence increase occurred during intromission and ejaculation (Fig. 8C,D). These results indicate that DA in the NAc is acutely released during episodes of sexual behaviors and may carry information regarding specific features of courtship and/or mating.

**Figure 8:**
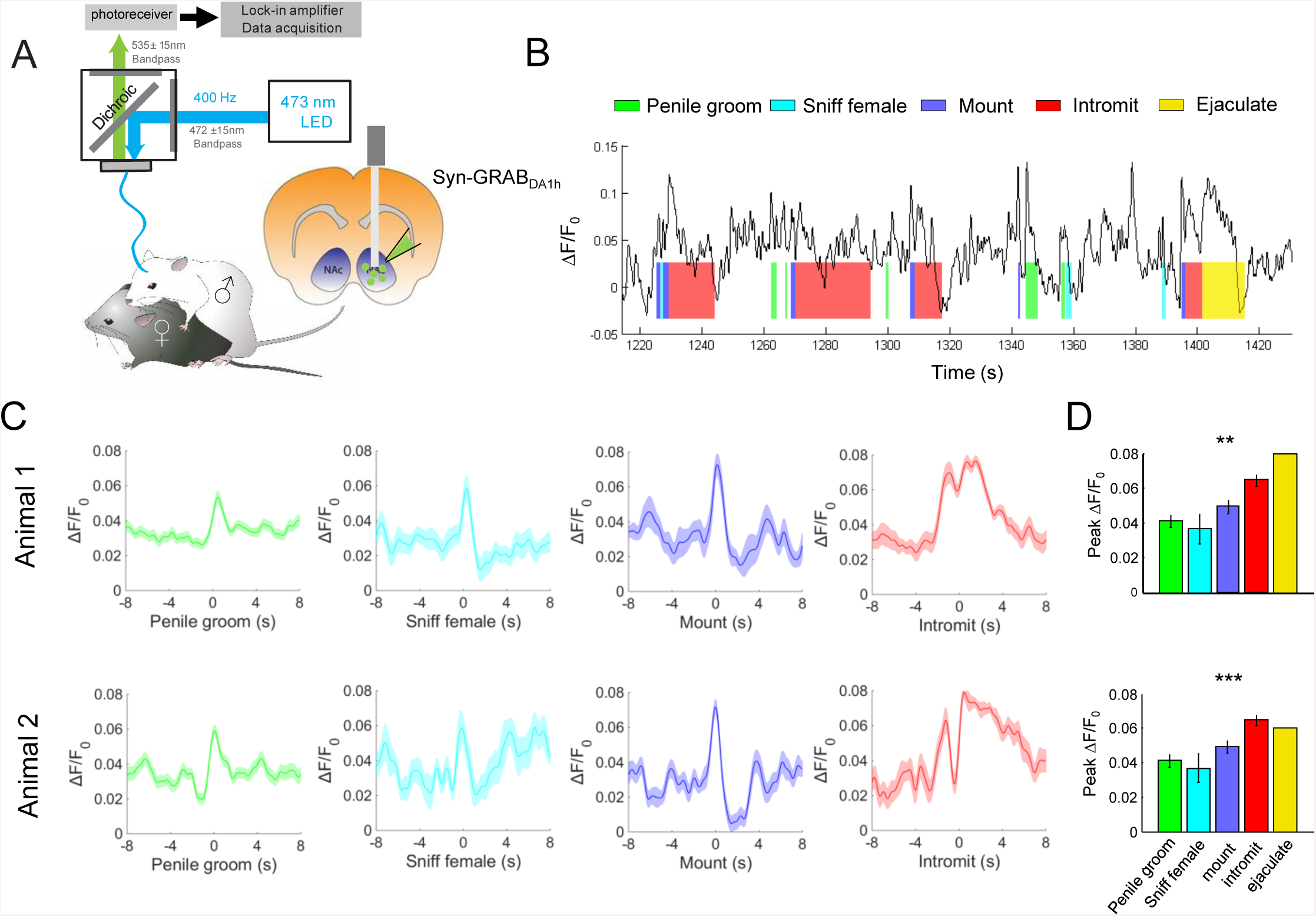
Acute DA release in the NAc measured during male sexual behaviors. (A) Schematic diagram showing the experimental design used to record GRAB_DA1h_ signals in the NAc of male mice during sexual behaviors. (B) Representative fluorescence changes in male mice during the indicated sexual behaviors. The shaded areas with colors indicate different behavioral events. (C) Post-event histogram (PETH) showing the GRAB_DA1h_ signals aligned to the indicated behavioral events. (D) Group data summarizing the average peak ΔF/F_0_ (baseline adjusted) during different indicated behaviors for the two mice shown in (C) (*n =* 2 mice; *p* = 0.007 for animal 1; *p* < 0.001 for animal 2). Error bars, ± SEM. One-way ANOVA performed; ***, *p* < 0.001; **, *p* < 0.01.

## Discussion

Here we describe the development and characterization of a pair of novel genetically-encoded sensors that enable specific, real-time detection of endogenous DA dynamics in various *ex vivo* and *in vivo* preparations and model systems. In acute mouse brain slices, GRAB_DA_ sensors were well suited to monitor stimulus-evoked DA release in mesolimbic pathway. In flies, GRAB_DA_ sensors were sufficiently sensitive to detect DA release in the MB triggered by odorants presented at physiologically relevant concentrations, and could also readily resolve DA release evoked by a single electrical stimulus. In transgenic zebrafish, GRAB_DA_ sensors were able to report DA release in the optic tectum in response to the chemogenetic activation of pretectal DANs. In mice, combining optogenetic stimulation with GRAB_DA_ sensors enabled the simultaneous optical manipulation and detection of DA signals *in vivo*. Finally, GRAB_DA_ sensors revealed real-time DA dynamics in the NAc of freely behaving mice as they underwent Pavlovian conditioning or engaged in sexual behaviors.

Compared to current methods used to measure DA release, our GRAB_DA_ sensors described here exhibit several clear advantages. First, GRAB_DA_ sensors are genetically-encoded by relatively small genes (∼2 kb), making them highly amenable to transgenic approaches and viral packaging. Second, GRAB_DA_ sensors have high sensitivity to DA. In response to DA, GRAB_DA1m_ and GRAB_DA1h_ sensors are capable of achieving maximal ΔF/F ∼ 90% with ∼10 nM and ∼100 nM affinities, respectively. In contrast, conventional GPCR-based FRET probes for detecting neurotransmitters are usually limited to a maximum FRET signal changes of ∼ 5% under optimal conditions, and less than that *in vivo* (Vilardaga et al., 2003; Ziegler et al., 2011). Third, GRAB_DA_ sensors have high specificity to DA: A range of experimental approaches, including application of multiple neurotransmitters and D_2_R antagonists, perturbation of DATs, or manipulation of DA synthesis pathways unequivocally support the molecular specificity of GRAB_DA_ sensors for DA. Notably, similar to the human D_2_R upon which they are based, both GRAB_DA_ sensors have a 10-fold higher affinity for DA than for NE (Lanau et al., 1997b); in contrast, FSCV is unable to discriminate between these two catecholamines (Fox and Wightman, 2016; Park et al., 2009). Finally, GRAB_DA_ sensors have extremely rapid response kinetics: GRAB_DA_ sensors report increases in DA levels with a rise time of ≤ 100 ms (Fig. 1G, 3D and 4I). Although this response time of GRAB_DA_ sensors is slower than voltammetry methods, it is still sufficiently rapid for reporting physiologically relevant DA dynamics and share response kinetics similar to WT GPCRs (Lohse et al., 2008), providing an accurate readout of their activation.

Because GRAB_DA_ sensors were engineered using the D_2_R, a G_i_-coupled GPCR, a potential concern is that overexpressing the GRAB_DA_ sensors may inadvertently activate pathways downstream of D_2_R. However, several lines of evidence argue against this possibility, including the negligible coupling between GRAB_DA_ sensors and both G protein-dependent and arrestin-dependent intracellular signaling pathways, alleviating this concern (Fig. 1I, 1J). This lack of coupling is presumably due to the steric hindrance imposed by the bulky cpEGFP moiety that replaces parts of the ICL3, which is the critical position for G protein or arrestin to interact with the GPCR (Luttrell and Lefkowitz, 2002; Neves et al., 2002). Consistent with minimal coupling between GRAB_DA_ sensors and downstream signaling pathways, *in vivo* Ca2+ imaging experiments using the Ca^2+^ sensor jRCaMP1a revealed no measurable alteration in Ca^2+^ signaling in neurons of transgenic flies that overexpressing GRAB_DA_ sensors (see Fig. S5).

In flies, DA signaling is critical for olfactory learning (Burke et al., 2012; Heisenberg, 2003; Liu et al., 2012; Schwaerzel et al., 2003). Here show that GRAB_DA_ sensors can be targeted to specific cells in the fly brain and used to probe odor-evoked DA dynamics in the MB of the living fly in real-time. It is reported that different types of DANs innervate different compartments of MBs, and these different dopaminergic pathways may play distinct roles in appetitive and aversive olfactory-dependent behaviors (Aso and Rubin, 2016; Cognigni et al., 2017). The GRAB_DA_ sensors developed here create new opportunities for exploring how distinct DA dynamics can correspond with specific compartments in the MB of the intact fly, particularly as the animal engages in different behavioral paradigms. Experiments in flies also illustrate the power of GRAB_DA_ sensors to probe DAT function *in vivo* by directly measuring extracellular DA level in real time. Similarly, we showed that GRAB_DA_ sensors readily respond to DA transients in the intact brain of the zebrafish larvae, providing a robust and convenient tool to examine DA dynamics in this classic vertebrate model system.

In addition to the morphological distinctions that can be made between various DANs in the mammalian CNS, breakthroughs in single-cell sequencing have further divided these neurons into a wide variety of cell types with distinct molecular features, suggesting high functional heterogeneity (Nair-Roberts et al., 2008; Ungless and Grace, 2012). Therefore, genetically-encoded GRAB_DA_ sensors provide a novel tool to explore patterns of DA release from genetically distinct DAN types, thereby facilitating the understanding of how they may be functionally specialized for different physiological and behavioral processes (Lammel et al., 2014; Pignatelli and Bonci, 2015). In fact, GRAB_DA_ sensors applied in the mouse faithfully reported the expected bi-directional regulation of DA levels in the NAc during different forms of Pavlovian conditioning. Consistent with the notion that DA can predict an anticipated reward (Schultz et al., 1997), the GRAB_DA_ sensor detected a robust, phasic increase in DA levels that shifted from reward onset to cue onset over the course of reward learning. Conversely, in naïve animals, a transient decrease in DA levels was triggered by the air puff onset and this decrease shifted to cue onset during aversive conditioning. An important goal of future studies will be to use GRAB_DA_ sensors to determine whether the same or different types of DANs are involved in appetitive vs. aversive learning.

Our experiments in freely behaving male mice provide new perspectives with respect to DA dynamics during sexual behaviors. Contrary to traditional views, GRAB_DA_ sensors expressed in the NAc revealed striking time-locked DA elevation immediately prior to and peak at the onset of various distinct sexual behaviors, consistent with a model where DA encodes behavioral motivation, anticipation, or arousal. These rapid changes in DA levels are in contrast with previous dialysis studies showing that DA levels in the NAc slowly increase during sexual behaviors, a difference that likely relates to the slow readout associated with dialysis-based methods. The targeted expression of GRAB_DA_ sensors could therefore provide a critical window into the coding strategy of DA release in complex behaviors. Moreover, because GRAB_DA_ sensors readily discriminate between DA and NE, they may be useful in studying cortical and subcortical regions in which dopaminergic and adrenergic inputs are intertwined.

Given that the crystal structure of the D_2_R was recently solved (Wang et al., 2018), future efforts can use this structural information to further tune the affinity, enhance the selectivity, and increase the signal-to-noise ratio in the next-generation of GRAB_DA_ sensors. Moreover, by adding a red fluorescent protein, GRAB_DA_ sensors can be readily transformed into ratiometric indicators, which could prove useful for more quantitative measurements of DA release across different experiments and preparations. Finally, a GPCR-based strategy was recently used to develop a genetically-encoded sensor (GACh) with high sensitivity and high selectivity for acetylcholine (ACh) (Miao Jing et al., 2018). Although the GACh sensor differs from the GRAB_DA_ sensors described here in that is based on the muscarinic Gq-coupled GPCR receptor M_3_R and an ICL3 loop derived from a Gs-coupled beta adrenergic receptor (Levitzki, 1988; Rasmussen et al., 2011), a feature common to this sensor is that the conformational changes in a GPCR induced by ligand binding are successfully harnessed and converted into a sizable increase in cpEGFP florescence. Given the diverse ligand-specificity of different GPCRs, a future goal will be to explore whether this principle can be expanded even further in order to develop sensors for the entire range of neurotransmitters and neuromodulators.

## Methods

### Animals

Wild-type Sprague-Dawley rat pups (P0) were used to prepare cultured cortical neurons. Wild-type C57BL/6 and TH-Cre mice (B6.FVB(Cg)-Tg(Th-cre)FI172Gsat/Mmucd obtained from MMRRC) were used to prepare the acute brain slices and *in vivo* experiments. All animals were maintained in the animal facilities and were family-or pair-housed in a temperature-controlled room with a 12-h/12-h light/dark cycle. All procedures for animal surgery and maintenance were performed using protocols that were approved by the Animal Care & Use Committees at Peking University, Chinese Academy of Sciences (CAS), New York University, University of California, San Francisco, and US National Institutes of Health, and were performed in accordance with the guidelines established by US National Institutes of Health guidelines.

To generate transgenic zebrafish, plasmids containing pTol2-elval3:GRAB_DA1m_ (25 ng/μL) and Tol2 mRNA (25 ng/μL) were co-injected into fertilized eggs, and founders were screened three months later. Transgenic zebrafish adults and larvae were maintained at 28 °C on a 14-h/10-h light/dark cycle.

To generate transgenic flies, the coding sequence of GRAB_DA1m_ was integrated into the pUAST vector using Gibson Assembly (Gibson et al., 2009), which was then used in P-element-mediated random insertion. Transgenic *Drosophila* lines carrying GRAB_DA1m_ on the chromosomes 2 and 3 with the strongest expression level after crossing with TH-GAL4 were used. The coding sequence of GRAB_DA1m-mut_ was incorporated into pJFRC28 (Pfeiffer et al., 2012) (Addgene plasmid #36431) using Gibson Assembly, and this plasmid was used to generate transgenic flies using PhiC31-mediated site-directed integration into attp40. The embryo injections were performed at Core Facility of Drosophila Resource and Technology, Shanghai Institute of Biochemistry and Cell Biology, CAS. Transgenic flies were raised on conventional corn meal at 25°C, with ∼70% humidity, under 12:12-h light-dark cycle.

The following *Drosophila* lines used in this study:

TH-GAL4, a gift and unpublished line generated by appending 2A-GAL4 to the last exon of TH, from Yi Rao, Peking University. C305a-GAL4 and 30y-GAL4, also gifts from Yi Rao. DTH^FS+/-^ple^2^/TM6B (Cichewicz et al., 2017), a gift from Jay Hirsh, University of Virginia. UAS-DAT-RNAi (TH01470.N), from Tsinghua Fly center, Tsinghua University. UAS-jRCaMP1a (Bloomington #63792), a gift from Chuan Zhou, Institute of Zoology, Chinese Academy of Sciences.

The following genotypes were used in the following figures: Fig. 4A-C

UAS-GRAB_DA1m_/cyo; TH-GAL4 (DANs)/TM6B

UAS-GRAB_DA1m-mut_/+; TH-GAL4/+

c305a-GAL4 (α’ and β’ Kenyon cells)/UAS-GRAB_DA1m_; DTHFS+/-ple2/+ (WT group)

c305a-GAL4/UAS-GRAB_DA1m_; DTHFS+/-ple^2^ (TH-deficient group)

Fig. 4D-K

UAS-GRAB_DA1m_/cyo; TH-GAL4/TM6B

UAS-GRAB_DA1m-mut_/+; TH-GAL4/+

Fig. 4L-O

UAS-GRAB_DA1m_/cyo; TH-GAL4/TM6B

UAS-GRAB _DA1m_/+; TH-GAL4/UAS-DAT-RNAi

Fig. S5

TH-GAL4/ UAS-jRCaMP1a

UAS-GRAB _DA1m_/+; TH-GAL4/UAS-jRCaMP1a

30y-GAL4/ UAS-jRCaMP1a

UAS-GRAB _DA1m_/+; 30y-GAL4/UAS-jRCaMP1a

### Molecular biology

Plasmids were generated using Gibson Assembly. DNA fragments were generated using PCR amplification with primers (Thermo Fisher Scientific) with 30-bp overlap. The fragments were assembled using T5-exonuclease (New England Biolabs), Phusion DNA polymerase (Thermo Fisher Scientific), and Taq ligase (iCloning). All sequences were verified using Sanger sequencing (Sequencing platform in the School of Life Sciences of Peking University). DNA encoding the various DA receptor subtypes (D_1_R-D_5_R) was generated using PCR amplification of the full-length human GPCR cDNAs (hORFeome database 8.1). For characterization in HEK293T cells, the GRAB_DA_ constructs were cloned into the pDisplay vector (Invitrogen), with an IgK leader sequence inserted upstream of the coding region. The IRES-mCherry gene was attached downstream of GRAB_DA_ and was used as a reference of membrane marker to calibrate of the signal intensity. Site-directed mutagenesis of the N- and C-terminal linker sequences in cpEGFP was performed using primers containing randomized NNB codons (48 codons in total, encoding the 20 possible amino acids; Thermo Fisher Scientific). Site-directed mutagenesis of the D_2_R gene was performed using primers containing the target sites. For the characterization in cultured neurons, the GRAB_DA_ constructs were cloned into the pAAV vector under the TRE promoter or the human synapsin promoter. The marker constructs RFP(mScarlet)-CAAX, EGFP-CAAX, KDELR1-EGFP, PSD95-mScarlet and synaptophysin-mScarlet were cloned into pEGFP-N3 vector.

### Expression of GRAB_DA_ in cultured cells and *in vivo*

HEK293T cells were cultured in DMEM supplemented with 10% (v/v) FBS (Gibco) and 1% penicillin-streptomycin (Gibco) at 37°C in 5% CO_2_. The cells were plated on 12-mm glass coverslips in 24-well plates and grown to ∼50% confluence for transfection. Transfection was performed by incubating the HEK293T cells with a mixture containing 1 μg of DNA and 3 μg of PEI for 6 h. Imaging was performed 24-48 h after transfection.

Rat cortical neurons were prepared from postnatal 0-day old (P0) Sprague-Dawley rat pups as previously described (Zhang et al., 2009). In brief, the cortical neurons were dissociated from the dissected rat brains in 0.25% Trypsin-EDTA (Gibco), and plated on 12-mm glass coverslips coated with poly-D-lysine (Sigma-Aldrich) in neurobasal medium containing 2% B-27 supplement, 1% GlutaMax, and 1% penicillin-streptomycin (Gibco). The cells were transfected 7-9 days later using the calcium phosphate transfection method. Imaging was performed 48-72 h after transfection.

For *in vivo* expression, Wild-type C57/BL6 mice with the age of P42-60 were first anesthetized by 2,2,2-Tribromoethanol (Avetin, 500 mg/kg) through intraperitoneal injection, or by isoflurane (RWD Life Science), and then placed in a stereotaxic frame to inject AAVs of GRAB_DA_ sensors into NAc with a microsyringe pump (Nanoliter 2000 Injector, WPI), or a microinjection pipette injector (Nanoject II, Drummond Scientific). The coordination of NAc was set as AP: −1.40 mm from Bregma, ML: 1.00 mm from the midline, DV: 3.90 mm from the brain surface. The injection was made unilateral with ∼300-500 nL per animal. In dual-color optical recordings experiments, the AAVs of hsyn-GRAB_DA1m_ and hsyn-tdTomato were injected in the dorsal striatum (AP = - 0.5mm, ML = ±2.5mm from bregma, and DV = −2.2mm from the brain surface), and AAV of Ef1a-DIO-C1V1-YFP was injected in the substantia nigra pars compacta (SNc) (AP = −3.1mm, ML = ±1.5mm from bregma, and DV = −4.0mm from the brain surface) in TH-Cre mice.

### Fluorescence imaging of HEK293T cells and cultured neurons

GRAB_DA_-expressing HEK293T cells and cultured neurons were imaged using an inverted Ti-E A1 confocal microscope (Nikon) and the Opera Phenix high content screening system (PerkinElmer). The Nikon confocal microscope was equipped with a 40×/1.35 NA oil immersion objective, a 488-nm laser and a 561-nm laser. During imaging, the HEK293T cells and cultured neurons were bathed or perfused in a chamber with Tyrode’s solution containing (in mM): 150 NaCl, 4 KCl, 2 MgCl_2_, 2 CaCl_2_, 10 HEPES and 10 glucose (pH 7.4). Solutions containing the drug/compound of interest (e.g., DA, Halo, 5-HT, histamine, NE, or ACh) were delivered via a custom-made perfusion system or via bath application. The chamber was fully cleaned with Tyrode’s solution and 75% ethanol between experiments. The GFP signals (e.g., the GRAB_DA_ sensors, the iGluSnFR or EGFP) were recorded using a 525/50 nm emission filter, and the RFP signals were collected using a 595/50 nm emission filter. The photostabilities of GRAB_DA1m_, GRAB_DA1h_, EGFP, and iGluSnFR were measured using 350-μW 488-nm laser illumination. Photobleaching was applied to the entire sensor-expressing HEK293T cell. The Opera Phenix high content screening system was equipped with a 60×/ 1.15 NA water immersion objective, a 488-nm laser, and a 561-m laser. The GRAB_DA_ signals were collected using a 525/50 nm emission filter, and the mCherry signals were collected using a 600/30 nm emission filter. Where indicated, the culture medium was replaced with 100 μl of Tyrode’s solution containing various concentrations of the indicated drug/compound. The fluorescence intensities of the GRAB_DA_ sensors were calibrated using mCherry as the reference.

### Fluorescence imaging of GRAB_DA_ in brain slices

Two weeks after the virus injection, the animals were anesthetized with Avetin and then decapitated. The brains were removed immediately and placed directly in cold slicing buffer containing (in mM): 110 choline-Cl, 2.5 KCl, 1.25 NaH_2_PO_4_, 25 NaHCO_3_, 7 MgCl_2_, 25 glucose, and 2 CaCl_2_. The brains were then sectioned into 200-μm thick slices using a VT1200 vibratome (Leica, Germany), and the sections were transferred into the oxygenated Ringer’s buffer containing (in mM): 125 NaCl, 2.5 KCl, 1.25 NaH_2_PO_4_, 25 NaHCO_3_, 1.3 MgCl_2_, 25 glucose, and 2 CaCl_2_; the slices were then allowed to recover in 34 °C for at least 40 minutes. For fluorescence imaging, the slices were transferred to an imaging chamber in an Olympus FV1000MPE two-photon microscope equipped with a 40×/0.80 NA water-immersion objective and a mode-locked Mai Tai Ti:Sapphire laser (Spectra-Physics) tuned to 920 nm for the excitation of GRAB_DA_ sensors and a 495∼540 nm filter for signal collection. For electrical stimulation, a concentric electrode (model #CBAEC75, FHC) was positioned near the NAc core under the fluorescence guidance, and the imaging and stimulation were synchronized using an Arduino board with custom programs. The stimulation voltage was set at 5-6 V, and the duration of each stimulation was set at 2 ms.

For immunostaining of brain sections, GRAB_DA_-expressing mice were anesthetized with Avetin, and the heart was perfused with 0.9% NaCl followed with 4% paraformaldehyde (PFA). The brain was then removed and placed in 4% PFA for 4 hours, then cryoprotected in 30% (w/v) sucrose for 24 hours. The brain was embedded into tissue-freezing medium, and 50-μm-thick coronal sections were cut using a Leica CM1900 cryostat (Leica, Germany). To label dopaminergic terminals and GRAB_DA_ in the NAc, tissue sections were rinsed and then immunostained with rabbit anti-TH antibody (1:100, Millipore, #ab152) and chicken anti-GFP antibody (1:500, Abcam, #ab13970), followed by an Alexa-555-conjugated goat-anti-rabbit and Alexa-488-conjugated goat-anti-chicken secondary antibodies. The immunostained tissue sections were imaged using the same Nikon confocal microscope in cell imaging.

### Fluorescence imaging of transgenic flies

Adult *Drosophila* (within 3 weeks of eclosion) were used for imaging experiments. The mounting and dissection protocols were as previously described (Liang et al., 2013). In brief, a section of rectangular cuticle between the eyes was removed to expose the brain, which was then bathed in saline, so called adult hemolymph-like solution (AHLS). The same Olympus two-photon microscope used for brain slices imaging was also used here. For GRAB_DA_ sensors imaging, 920-nm excitation laser and 495∼540-nm filter were used. For jRCaMP1a, 1000-nm excitation laser and 575∼630-nm filter were used. For olfactory stimulation, the odorant isoamyl acetate (Sigma-Aldrich; Cat. #306967) was firstly diluted 200-fold in mineral oil in a bottle; this dilution was subsequently diluted 5-fold in air, which was then delivered to the fly’s antenna at a rate of 1000 ml/min. Compounds such as Halo and cocaine were added directly to the AHLS to their final concentration, and the following experiments were performed 10 min after compound application. For electrical stimulation, a glass electrode (resistance ∼0.2 MO) was placed in the region of the DANs in the MB and the stimulation voltage was set at 20∼80 V. Arduino was used to synchronized stimulation delivery and imaging with custom code. The sampling rates during olfactory stimulation and electrical stimulation was 2.7 Hz and 12 Hz, respectively.

### Fluorescence imaging and chemogenetics in zebrafish

All experiments were performed on 5 days-post-fertilization (5 dpf) larvae in 10% Hank’s solution containing (in mM): 140 NaCl, 5.4 KCl, 0.25 Na_2_HPO_4_, 0.44 KH_2_PO_4_, 1.3 CaCl_2_, 1.0 MgSO_4_, and 4.2 NaHCO_3_ (pH 7.2). Imaging of Tg (elval3:GRAB_DA1m_/DAT:TRPV1-TagRFP) larvae at 5 dpf was performed with an inverted confocal microscope (Olympus FV3000, Japan) using a 30X oil-immersion objective (1.05 N.A., morphology imaging) or an upright confocal microscope (Olympus FV1000, Japan) using 40X water-immersion objective (0.8 NA, time-lapse imaging). After the larvae were paralyzed with a-bungarotoxin (100?μg/ml, Sigma), they were mounted dorsal side up in 1.5% low melting-point agarose (Sigma) and then immersed in an extracellular solution consisting of (in mM): 134 NaCl, 2.9 KCl, 4 CaCl_2_, 10 HEPES and 10 glucose (290?mOsmol/L, pH 7.8). For imaging the morphology, images were acquired with a field of view consisting of 1,024 pixels × 1,024 pixels with spatial resolution of 0.414 × 0.414 × 1 μm^3^ (x × y × z). For bath application of compounds, dopamine (100 μM in 1 mM ascorbic acid solution, Sigma) was added by pipette at ∼ 4 min and haloperidol (50 μM in DMSO, Tocris) at ∼12 min. These images were acquired with a view field of 640 × 640 pixels with spatial resolution of 0.497 × 0.497 μm^2^ (x × y) at ∼1.5 Hz. For functional imaging, small anterior dissections initiated in ventricles were made, after which a glass pipette containing the TRPV1 agonist capsaicin (50 μM in absolute ethanol, Tocris) was advanced through the incision and placed near the cell bodies of the DANs. To activate the DANs, 5 pulses of puffs (9-10 psi, 100-ms) were delivered with 1 min interval. The larvae were bath in Halo (50 μM in DMSO, Tocris) for 10 min before imaging. These images were acquired with a field of view consisting of 800 × 800 pixels with spatial resolution of 0.397 × 0.397 μm^2^ (x × y) at ∼1 Hz.

### Fiber Photometry recording in freely moving mice

In all-optic experiments in Fig. 6, optical fiber probes (105 μm core /125 μm cladding) were implanted in the dorsal striatum and in SNc 4 weeks after the virus injection. Fiber photometry recording in the dorsal striatum was performed using a 50-μW 470-nm LED, and C1V1 in the SNc was stimulated using a 9.9- mW 561-nm laser. The measured emission spectra of GRAB_DA1m_ and tdTomato were fitted using a linear unmixing algorithm https://www.niehs.nih.gov/research/atniehs/labs/ln/pi/iv/tools/index.cfm. The coefficients of GRAB_DA1m_ and tdTomato generated by the unmixing algorithm were used to represent the fluorescence intensities of GRAB_DA1m_ and tdTomato, respectively. To evoke C1V1-mediated DA release in the dorsal lateral striatum, pulse trains (10-ms pulses at 10 Hz for 1 s) were delivered to the SNc using a 9.9-mW, 561-nm laser. In other experiments in Fig. 7 and 8, an optic fiber (Thorlabs, FT200UMT,FT400UMT or BFH48-400) was attached to the implanted ferrule (Thorlabs, SF440-10) via a ceramic sleeve. A 400-Hz sinusoidal blue LED light (30 μW) (LED light: M470F1; LED driver: LEDD1B; both from Thorlabs) was bandpass filtered (passing band: 472 ± 15 nm, Semrock, FF02-472/30-25 in Fig.8 or 460-490nm in Fig.7) and delivered to the brain to excite GRAB_DA_. The emission light then traveled through the same optic fiber, was bandpass filtered (passing band: 534 ± 25 nm, Semrock, FF01-535/50 in Fig.8 or 500-550nm in Fig.7), detected by a Femtowatt Silicon Photoreceiver (Newport, 2151) and recorded using a real-time processor (RZ5, TDT). The envelope of the 400-Hz signals that reflects the intensity of the fluorescence signals was extracted in real-time using a custom TDT program.

### Behaviors

For the auditory conditioning task, mice were recovered for >3 days after surgery, and then water-restricted until reaching 85-90% of its original body weight and then prepared for behavior training. In the first Pavlovian task, the mice were trained on two frequency modulated pure tone auditory cues of 500 ms in duration, centered around 2.5 kHz and 11 kHz. For each mouse, one of the two tones was pseudo-randomly assigned to be the reward-predictive tone. Reward (water sweetened with 10% sucrose) was delivered through a water spout in front of the mouth following the reward-predictive cue with a variable 500-1500 ms delay. Rewarded and unrewarded trials were randomly interleaved with a variable inter-trial interval of 8-20 s. Mice experienced 200 trials (∼100 rewards) per day in sessions lasting ∼45 min.

In the subsequent Pavlovian conditioning task, the mice were trained on an auditory conditioning task, in which three pairs of auditory cues → outcomes pairs (or CS-US pairs; 8 kHz pure tone → 9 μl water; white noise → brief air puff to the face; and 2 kHz pure tone → no response) were delivered at random with a 10 −20 s randomized inter-trial interval. The duration and intensity of each auditory cue was 1 s and 70dB, respectively. The respective outcomes was delivered 1 s after the end of each auditory cue. The behavioral setup consisted of a custom-built apparatus allowing head fixation of the mouse’s head to a Styrofoam rod (diameter: 15 cm). Rotation of the Styrofoam rod, which corresponds to the animal’s running speed, was detected using an optical rotatory encoder. Licking behavior was detected when the mouse’s tongue contacted the water delivery tube. Each lick signal was processed using an Arduino UNO board with custom code and sent digitally to the training program (written in MATLAB) via a serial port. Water delivery was precisely controlled using a stepping motor pump, and the air puff (15 psi, 25-ms duration) was controlled using a solenoid valve. Timing of the pump and valve was controlled using the same Arduino UNO board used for lick detection, which also provided synchronization between the training program and the data acquisition system (RZ2 processor, Tucker-Davis Technologies). During first two days of each training session, the outcomes were delivered without the prediction cues.

The sexual behaviors are defined following conventions in previous literature (Hull and Rodriguez-Manzo, 2009). In details, sniffing female was defined as the male’s nose coming in close proximity to the female’s facial, body, and/or urogenital areas. “Mount” was defined as when the male posed his forelegs over the female’s back and with his hindfeet on the ground accompanying shallow pelvic thrusts. The mounting onset was defined as the moment at which the male tried to clasp female back. “Intromission” was defined as a deep rhythmic thrust following mounting. The onset of intromission was defined as the time at which the male performed the first deep thrusting toward the female with vaginal penetration. “Penile grooming” was defined when a male animal repeated grooming for his urogenital area after intromission and ejaculation. Ejaculation is detected when the male stopped thrusting and freeze for seconds. The putative ejaculation event was confirmed by the presence of vaginal copulatory plug.

### Data analysis

For imaging experiments in HEK293T cells, neurons, acute brain slices and transgenic flies, images were first analyzed using Image J software (National Institutes of Health), and then analyzed using Origin 9.1 (OriginLab) and MATLAB (MathWorks) with custom-written scripts. The data in acute brain slices and flies were first binned by 2x and averaged to generate representative traces.

For fiber photometry data, the signal baseline was first obtained by the MATLAB function “msbackadj” with a moving window of 25% of the total recording duration in Fig.8, or by subtracting 2^nd^ order exponential fitted data from the raw data after 10.17 Hz binning in Fig. 7. The fluorescence responses were indicated by ΔF/F_0_ in Fig.8, which was calculated as (F_raw_ –F_baseline_)/F_baseline,_ or by Z score in Fig.7. To analyze event-evoked changes in DA release, we aligned each trial to the onset or offset of the behavior, and calculated the peri-stimulus time histogram (PSTH). To compare PSTH changes during different phases of the training, we used data from the 2^nd^ day as naive, the 5-10^th^ day as trained and >10^th^ day as well-trained, and normalized the PSTH of each animal by water-evoked response during early training. The peak response during a behavior was calculated as the maximum ΔF/F_0_ during the behavior minus the average ΔF/F_0_ in the duration-matched period 2s prior to the behavior onset in Fig.8. The response to the CS was defined as the peak of the normalized PSTH between the CS onset and the US onset, and the response to US was calculated similarly using data from the US onset to data collected 2 s after the US onset in Fig.7.

Except where indicated otherwise, group differences were analyzed using the Student’s *t*-test, test, sign-rank test, One-Way ANOVA or post-hoc Tukey’s test.

Except where indicated otherwise, all summary data presented as the mean ± SEM.

## Acknowledgements

This work was supported by the National Basic Research Program of China (973 Program; grant 2015CB856402), the General Program of National Natural Science Foundation of China (project 31671118 and project 31371442), an NIH brain initiative grant NS103558, and the Junior Thousand Talents Program of China to Y.Li and to S.Zhang. Additional support was provided by the Ministry of Science and Technology of the People’s Republic of China (2017YFA0505703) to M.Xu. We thank Y. Rao for sharing the 2-photon microscope and X. Lei for the platform support of Opera Phenix high content screening system at PKU-CLS. We thank the Sequencing platform in the School of Life Sciences of Peking University. We thank R. Mooney, Y. Huan and L. Luo for valuable feedback of the manuscript.

## Author Contributions

Y.L. conceived and supervised the project. F.S., M.J., and J.Z. performed experiments related to sensor development, optimization and characterization in culture HEK cells, culture neurons, brain slices and transgenic flies, with the initial work from YC. L. and J. F., and help from Z.Y. F.L. and J.D. designed and performed experiments on transgenic fish. J.Z., Y.G., T.Y., W.P., S.O., L.W., S.Z., D.L., M.X., A.K., and G.C. designed and performed experiments in behaving mice. All authors contributed to data interpretation and data analysis. Y.L. wrote the manuscript with input from F.S., J.Z., M.J., D.L., S.O., M.X. and help from other authors.

---

**Supplementary Figure S1:**
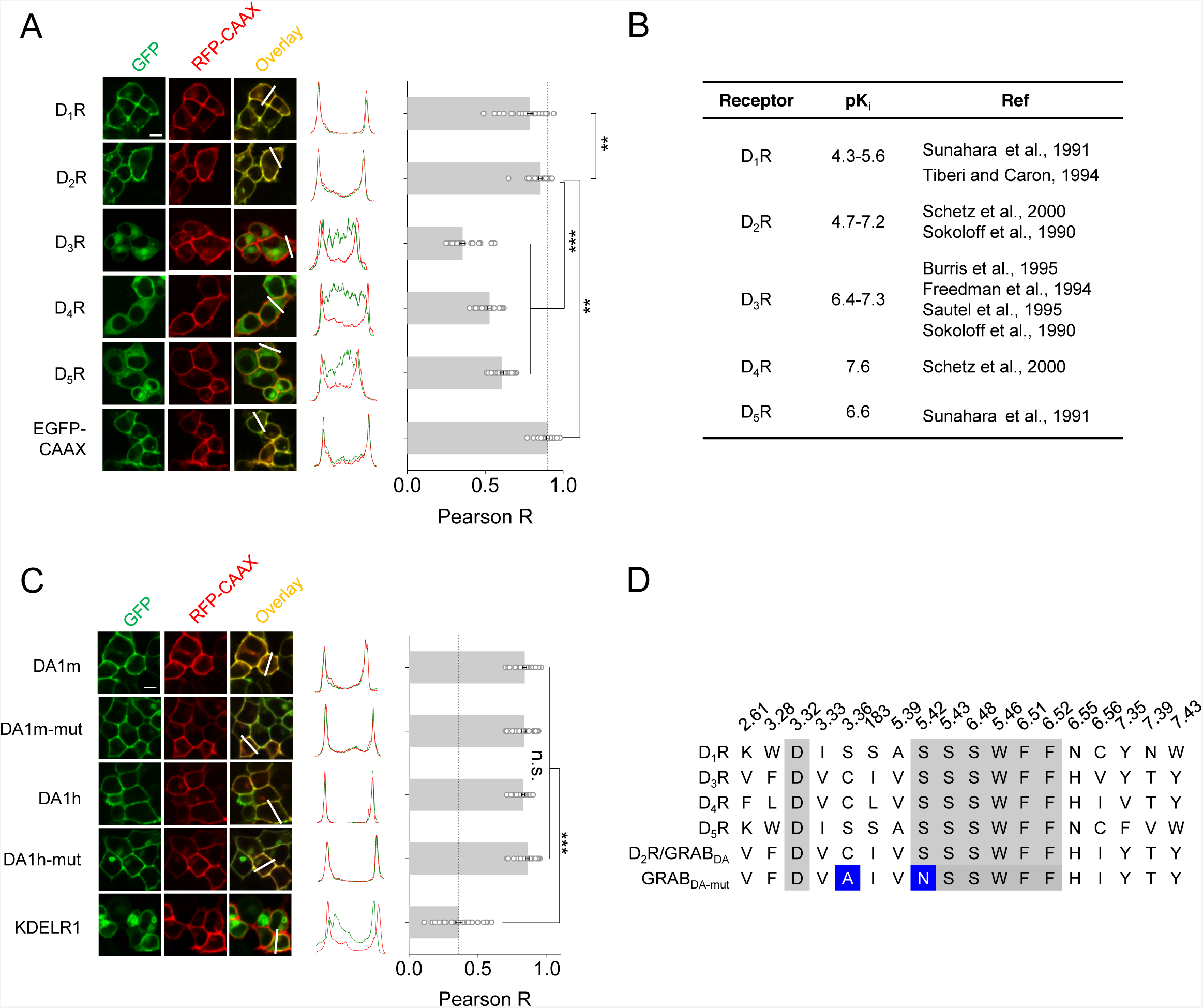
Comparison of DRs and DR-based chimeras, related to Fig. 1. (A) The fluorescence and membrane trafficking of all five DRs with cpEGFP insertion. A membrane localized RFP (RFP-CAAX) was co-expressed to indicate the plasma membrane and EGFP-CAAX was set as a control. Left, the fluorescence images of HEK293T cells expressing all five DR-based chimeras (green) and RFP-CAAX (red). Middle, the normalized line-scanning plots of the fluorescence signals in both green and red channels. Right, Pearson colocalization ratios between the DR-based chimeras and RFP-CAAX (*n =* 30/2 for each protein; *p* < 0.001 comparing D_2_R with D_3_R, D_4_R and D_5_R; *p* = 0.001 between D_2_R and EGFP-CAAX; *p* = 0.006 between D_2_R and D_1_R). (B) DA binding affinities of five subtypes of DRs. (C) Similar as (A), except different D_2_R based GRAB_DA_ sensors were characterized, including GRAB_DA1m_, GRAB_DA1m-mut_, GRAB_DA1h_, GRAB_DA1h-mut_ and Golgi marker KDELR1-GFP as a control (*n* = 30/2 for each protein; *p* > 0.05 among GRAB_DA_ sensors; *p* < 0.001 comparing KDELR1 with GRAB_DA_ sensors). (D) Sequence alignment of the binding pockets of D_1_R-D_5_R and D_2_R-based GRAB_DA_ sensors. Two blue-shaded amino acids indicate C118A and S193N mutation sites. Scale bars in (A) and (C), 10 μm.

**Supplementary Figure S2:**
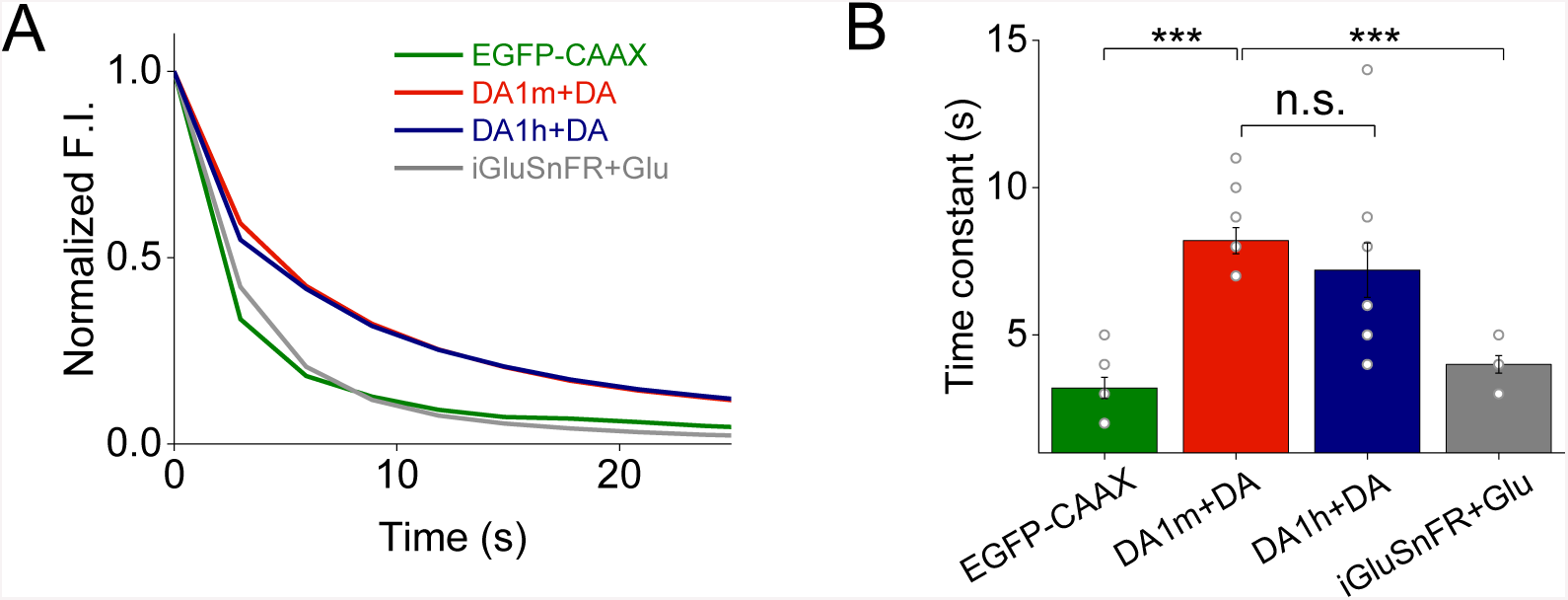
Photostability of GRAB_DA1m_ and GRAB_DA1h_ compared with other fluorescent probes, related to Fig. 1. (A) Representative photobleaching curves of GRAB_DA1m_, GRAB_DA1h_, EGFP-CAAX and iGluSnFR expressed in HEK293T cells under confocal imaging (488 nm laser with ∼350 μW intensity). (B) The group data of fluorescence decrease time constants of GRAB_DA1m_, GRAB_DA1h_, EGFP-CAAX and iGluSnFR (*n* = 10/3; *p* = 0.350 between GRAB_DA1m_ and GRAB_DA1h_; *p* < 0.001 comparing GRAB_DA1m_ with EGFP-CAAX and iGluSnFR).

**Supplementary Figure S3:**
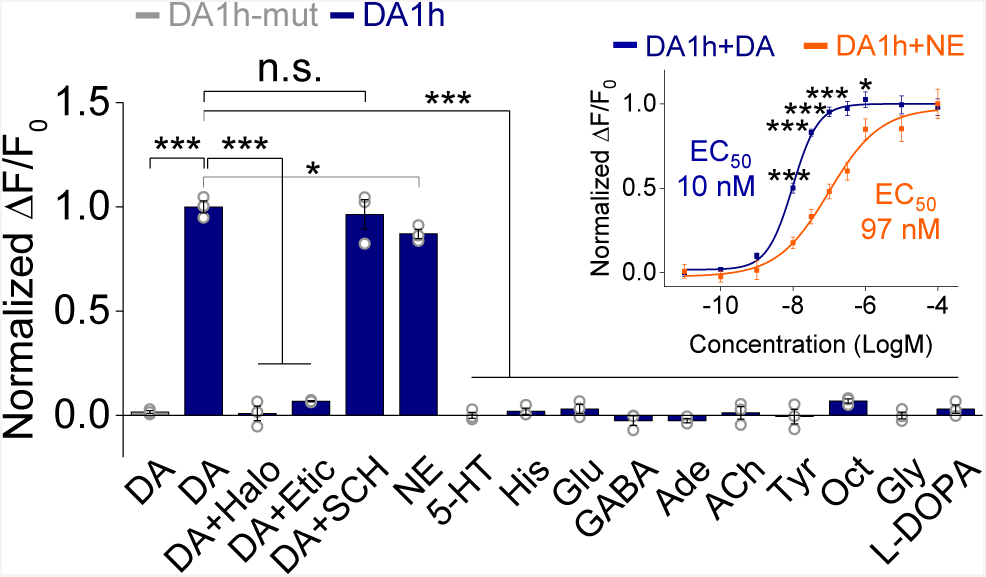
The selectivity of GRAB_DA1h,_ related to Fig. 1. Normalized fluorescence responses of GRAB_DA1h_- or GRAB_DA1h-mut_-expressing cells to the application of 1 μM different compounds, including: DA, DA with Halo, DA with Etic, DA with SCH, 5-HT, His, Glu, GABA, Ade, ACh, Tyr, Oct, Gly and L-Dopa (the first bar shows GRAB_DA1h-mut_-expressing cells in response to DA; *n* = 3 wells with each well of 200-400 cells; *p* < 0.001 for DA-induced responses between GRAB_DA1h_ and GRAB_DA1h-mut_; *p* = 0.66 for GRAB_DA1h_ responses induced by DA comparing with DA+SCH; *p* = 0.037 for GRAB_DA1h_ responses induced by DA comparing with NE; *p* < 0.001 for GRAB_DA1m_ responses induced by DA comparing with DA+Halo, DA+Etic, 5-HT, His, Glu, GABA, Ade, ACh, Tyr, Oct, Gly and L-DOPA). The inset, normalized dose-dependent fluorescence responses of GRAB_DA1h_-expressing cells to the application of DA (blue) or NE (orange) (*n* = 6 wells with each well of 200-400 cells; *p* < 0.001 at −8, −7.5, −7 and −6.5; *p* = 0.049 at −6).

**Supplementary Figure S4:**
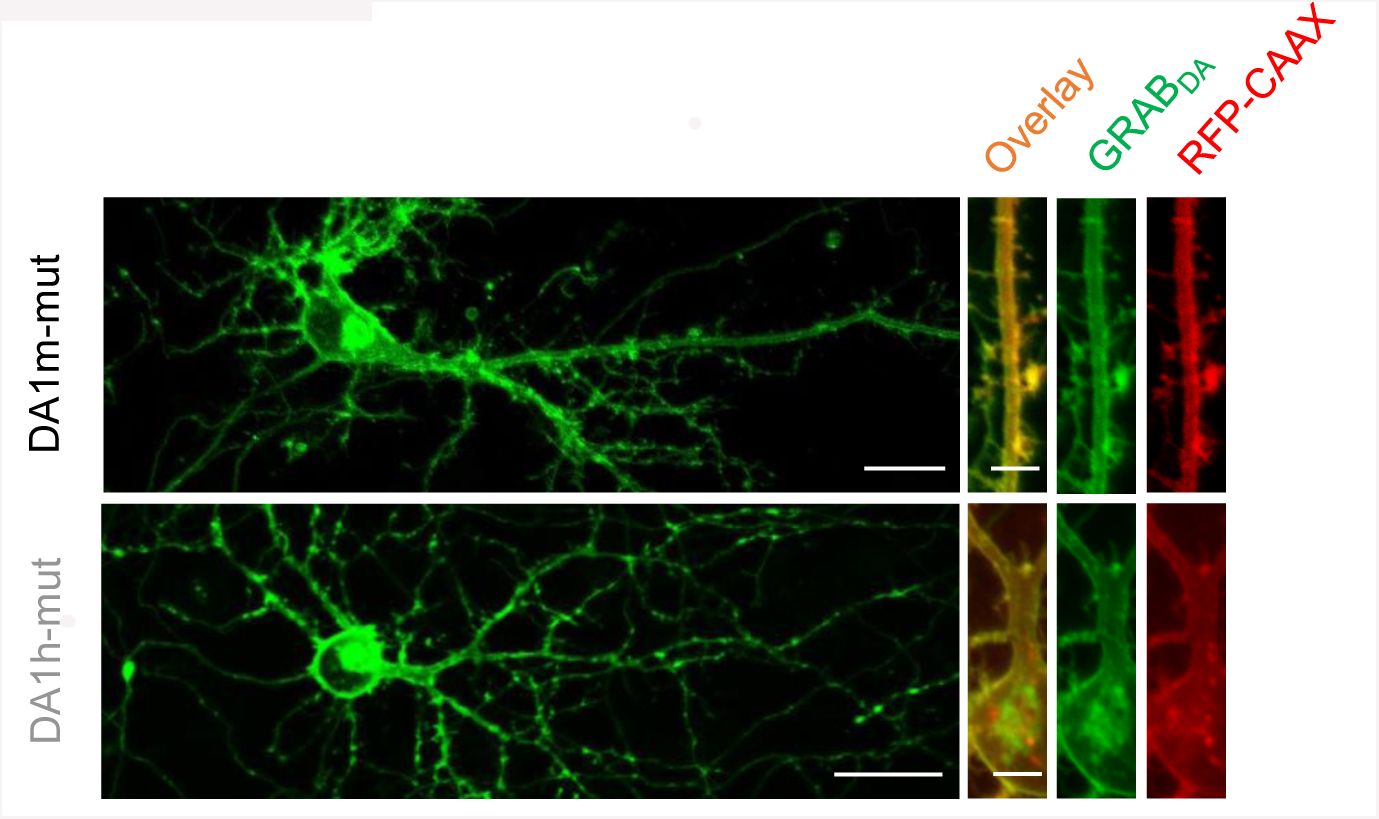
Expression and membrane trafficking of GRAB_DA1m-mut_ and GRAB_DA1h-mut_ in cultured neurons, related to Fig. 1. The fluorescence images of GRAB_DA1m-mut_- and GRAB_DA1h-mut_-expressing neurons under confocal microscope. RFP-CAAX was co-expressed in the same neuron to indicate the plasma membrane. Scale bars of whole views, 20 μm. Scale bars of zoom-in views, 5 μm.

**Supplementary Figure S5:**
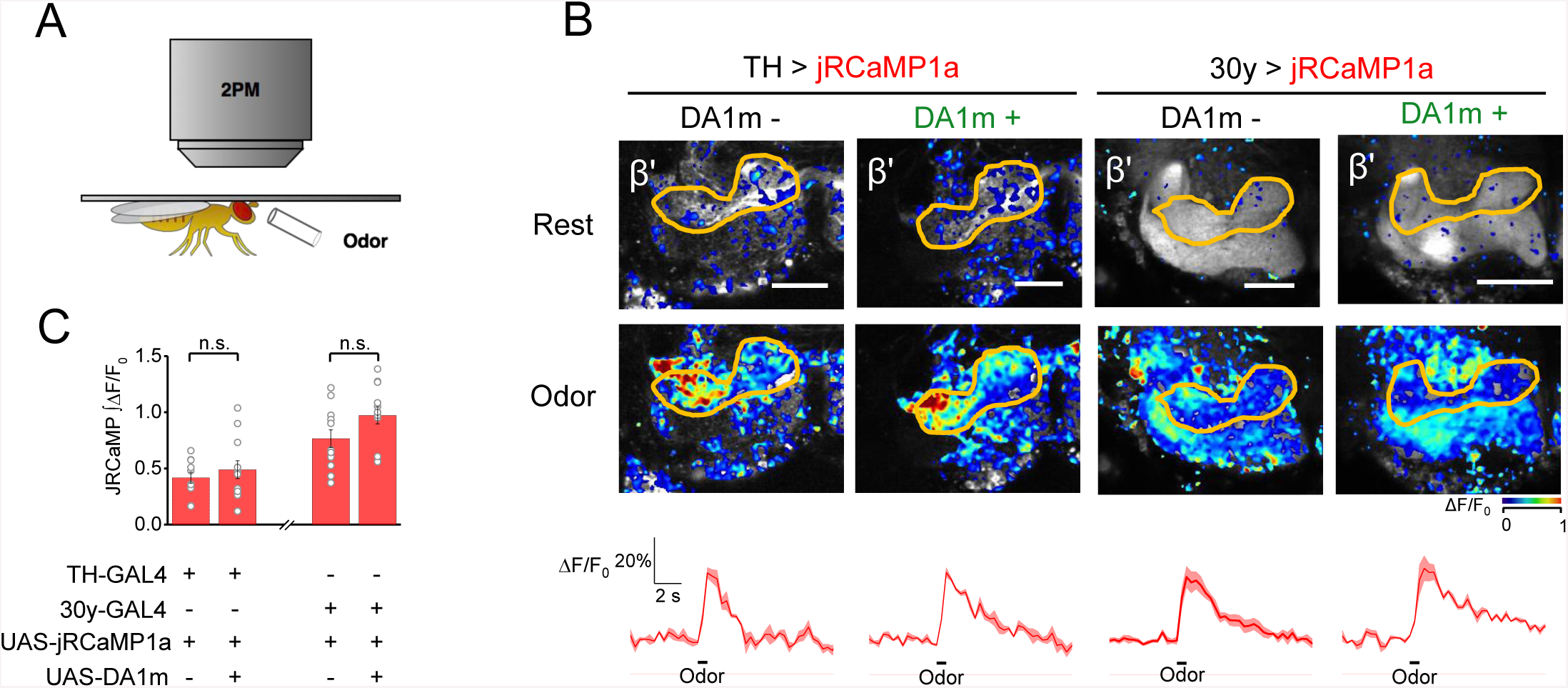
GRAB_DA1m_ expression has no effect on the odor-evoked Ca2+ signaling, related to Fig. 4. (A) The schematic of *in vivo* olfactory stimulation experiment under two-photon microscope. (B) Representative pseudo-color images and traces of fluorescence responses of jRCaMP1a-exprssing DANs (left) or Kenyon cells (right) to 1 s olfactory stimulation. Traces are 3-trial-averaged results from one fly, and are shaded with ± SEM. Scale bars, 25 μm. (C) The integrals of jRCaMP1a signals are summarized. Each dot represents data from one fly (TH > jRCaMP1a: *n =* 10 flies; TH > jRCaMP1a, GRAB_DA1m_: *n =* 11 flies; 30y > jRCaMP1a: 11 flies; 30y > jRCaMP1a, GRAB_DA1m_: *n =* 12 flies; *p* = 0.503 between TH > jRCaMP1a and TH > jRCaMP1a, GRAB_DA1m_; *p* = 0.097 between 30y > jRCaMP1a between 30y > jRCaMP1a, GRAB_DA1m_). Error bars, ± SEM. Student’s t-test is performed; n.s., not significant.

